# Fifty years of reduction in sulphur deposition drives recovery in soil pH and plant communities

**DOI:** 10.1101/2022.07.04.498654

**Authors:** Fiona M. Seaton, David A. Robinson, Don Monteith, Inma Lebron, Paul Bürkner, Sam Tomlinson, Bridget A. Emmett, Simon M. Smart

## Abstract

1. Sulphur deposition through rainfall has led to species loss and ecosystem degradation globally, and across Europe huge reductions in sulphur emissions since the 1970s were expected to promote the recovery of acidified ecosystem. However, the rate and ecological impact of recovery from acidification in terrestrial ecosystems is still unclear as is the influence of management and climate, as to date there has been no long-term spatially extensive evaluation of these changes.
2. Here we present data from thousands of sites across Great Britain surveyed repeatedly from 1978-2019 and assess change in soil pH and plant acidity preference (Ellenberg R) in response to atmospheric deposition of sulphur and nitrogen. We analyse change in grasslands managed for pasture, referred to as high-intensity habitats, and compare to semi-natural habitats comprising rough grassland, broadleaved woodland, bog and heathland, referred to as low-intensity habitats.
3. Soil pH increased from 1978 to 2007 but then decreased between 2007 and 2019, resulting in a net increase of ~0.2 pH units in low-intensity habitats but no change in high-intensity habitats. The community average Ellenberg R increased in semi-natural habitats by ~0.2 units but remained stable in intensive grasslands.
4. In semi-natural habitats, but not intensive grasslands, these changes in plant community composition were associated with the soil pH changes which were in turn linked to decreasing sulphur deposition and differences in rainfall.
5. Nitrogen deposition, which was relatively stable over the survey period, showed no additional effect upon soil acidity once sulphur deposition was accounted for.
6. **Synthesis:** Our results provide conclusive evidence that reductions in acid emissions are stimulating the gradual recovery of chronically acidified terrestrial ecosystems at a whole-country scale, while also suggesting this recovery is being compromised by changing climate and land management.

## Introduction

Atmospheric deposition of sulphur (S) and nitrogen (N) compounds has led to the widespread acidification of natural ecosystems and resulting loss of biodiversity and changes in ecosystem functions (Bytnerowicz et al., 2007; Kirk et al., 2010; Maskell et al., 2010). In Europe and North America S deposition dramatically declined in the late 20^th^ and early 21^st^ century due to switches away from fossil fuels and more effective monitoring and control of S emission sources. In in China more recent declines in emissions are only now beginning to result in decreases in precipitation acidity while in other countries such as India emissions are still increasing (Grennfelt et al., 2020; S. J. Smith et al., 2011; Zhang et al., 2020). This raises the possibility of being able to examine ecological recovery within areas now exposed to a long period of reduced S deposition in order to predict and manage change in areas where S deposition is still high. Within the UK there was a 96% reduction in sulphur dioxide (SO_2_) emissions from 1970 to 2019, concurrent with a 57% decline in emissions of oxides of nitrogen (NO_x_). Ammonia (NH_3_) emissions have remained relatively stable, showing only a 13% decrease from 1980-2019 (NAEI, 2019). The late 20^th^ century reduction in S deposition to natural ecosystems has been linked to decreases in soil and water acidity (Hughes et al., 2012; Kirk et al., 2010; Reynolds et al., 2013), while N deposition has been associated with eutrophication effects upon plant communities through increased accumulation of N compounds within soils (Bytnerowicz et al., 2007; Schweiger & Beierkuhnlein, 2017; Tipping et al., 2021).

Decreasing soil acidity should influence plant community composition towards species adapted to less acidic, more nutrient rich environments (Peppler-Lisbach et al., 2020; Stevens et al., 2011). Decreasing aluminium solubility and toxicity with increasing pH is likely to increase soil microbial activity and facilitate colonisation of acid-sensitive plant species (Jones et al., 2019; Zhao & Shen, 2018). In addition, increased macronutrient availability, associated with an increase in soil pH (Zhao & Shen, 2018) and reduction in ionic aluminium, could theoretically unlock the eutrophying potential of accumulated N, particularly in areas most exposed to N deposition historically (Schneider et al., 2018; Smart et al., 2014). Nitrogen deposition not only acts as a eutrophying agent but may also add to the acid load, depending on how it is processed within catchments. Over the late 20^th^ century, plant communities across Europe were observed to be shifting towards communities more adapted to higher nutrient conditions, which has largely been attributed to changes in management intensity and N deposition but could also be related to changes in acidity (Diekmann & Dupré, 1997; Duprè et al., 2010; Smart et al., 2003).

Changes in the atmospheric deposition of acidity and nutrient N have been occurring at a time of increasing pressures from changes in climate and land use, and the influence of interactions between these drivers on natural ecosystems have been identified (Bytnerowicz et al., 2007). Increasing temperature has been shown to influence plant and soil functional response to acid deposition in rainfall, in some cases leading to greater effects – e.g. increased susceptibility of plants to air pollution at higher temperatures due to increased stomatal opening – and in other cases leading to reduced effects – e.g. soil respiration increase with temperature being reduced by acid rain (Bytnerowicz et al., 2007; Chen et al., 2021). Changes in precipitation will influence both deposition of atmospheric pollutants and soil acidity directly through changes in soil moisture (Hole & Engardt, 2008; Marwanto et al., 2018; Yu et al., 2020). As soil moisture increases soil pH increases, due to increased anaerobic conditions which lead to a microbially mediated increase in proton consumption during denitrification reactions (Dobbie & Smith, 2001; Zárate-Valdez et al., 2006). Recent analysis of soil pH trends across England and Wales suggested that changes in management practices, and particularly the cessation of liming due to reduced subsidies, may have confounded recent soil pH recovery (Rawlins et al., 2017; Seaton et al., 2021). This shows that soils may in future either stabilise at the current pH, or potentially begin to acidify again with corresponding effects upon plant communities and ecosystem health and reversal of ecosystem recovery. On the other hand, recent proposals to increase carbon sequestration in soils by incorporating basalt or alternative rocks into cropland would lead to alkalinisation of the soil (Beerling et al., 2018). Understanding the interrelationship between changes in management, climate, soil pH and vegetation is necessary to better predict the consequences of these technologies upon biotic communities across realistic combinations of ecosystem and land-use.

Here we use data from the UKCEH Countryside Survey of Great Britain to evaluate trends in soil pH and plant community acidity preference and examine their connection to atmospheric S and N deposition. The Countryside Survey (CS) comprises a representative sample of co-located vegetation and soils data from the British landscape. It was first carried out in 1978 and since then in 1990, 1998, 2007 and most recently a partial resurvey starting in 2019 (Wood et al., 2017). This long-term, large scale monitoring enables us to better evaluate change in semi-natural ecosystems in response to a wide range of concurrent biotic and abiotic pressures and over timescales relevant to plant community assembly and related ecological processes. Due to the extensive quality assurance protocols that are part of this survey we are also able to incorporate quantitative assessment of measurement error in the soils and vegetation data into our Bayesian modelling to more robustly detect change over time and attribute it to various drivers. We selected sites that were within semi-natural habitats (low-intensity management) or intensively managed grassland (high-intensity management) habitats and estimated change over time in soil pH and vegetation community acidity preference, represented by Ellenberg R. Ellenberg R is an ordinal score assigned to plant species that ranges from one to nine and represents the preferred acidity of that plant species, which can be averaged over the plant community. It is part of a suite of Ellenberg scores originally developed for Central Europe and adapted for Great Britain by Hill et al. (2000). We hypothesised that, over the last five decades of major reductions in acid deposition, soil pH and Ellenberg R values would have increased due to recovery from acidification, but that high intensity management sites would show less change due to reducing acid deposition occurring at the same time as a reduction in liming which had artificially increased soil pH in the mid-20^th^ century. We compared measurements of Ellenberg R with and without weighting by cover estimates and from larger and smaller nested plot sizes, to evaluate whether the increased sensitivity of cover-weighted estimates was counteracted by increased measurement error. We then tested whether changes in soil pH were associated with decreases in atmospheric S deposition and with cumulative N deposition at each site reflecting legacy effects of acidifying N deposition in parallel with a eutrophying effect (Rowe et al., 2020). The relative influences of S deposition and N deposition upon pH change were evaluated within a Bayesian framework, also including the difference in field season rainfall between survey years to incorporate effects of changes in climate as represented by changing precipitation. We also hypothesised that the changes in soil pH resulted changes in Ellenberg R and hence plant species composition, and tested this using a multivariate Bayesian model.

## Methods

### Field survey

The data presented here is from the UKCEH Countryside Survey, a repeated programme of field surveys with locations spread across the GB countryside (Figure 1, Carey et al., 2008). We include data from 1978, 1990, 1998, and 2007 as well as results from the recent resurveys of the previously surveyed sites. The recently surveyed sites were mostly resurveyed in 2019, however a small number in Wales were surveyed in 2016 (Emmett & the GMEP team, 2017). Within each 1 km square there are multiple vegetation plots, including five square plots measuring 14.14 by 14.14m. These plots are surveyed for plant composition in a nested structure, with soil samples taken to 15 cm depth from a central position on the corner of the inner 2 by 2m nest within the plot. Within the 2019 data a subset of plots had only the inner 2 by 2m square surveyed for vegetation (221 plots out of 559 plots). All vegetation data is available at the NERC Environmental Information Data Centre (EIDC), see data availability statement. Broad habitats were assigned to each plot by the surveyors according to JNCC guidance (Jackson, 2000). These broad habitats were grouped into high intensity management and low intensity management land. The high intensity management land comprised improved and neutral grassland where the summed cover of agricultural forage species, i.e. *Lolium perenne, Lolium multiflorum* and *Trifolium repens*, is over 25%. The low intensity management land included the rest of the neutral grassland, acid grassland, broadleaved woodland, dwarf shrub heath, bog, bracken, fen, marsh, swamp, and calcareous grassland. Coniferous woodland, arable, coastal and urban habitats were not included as we expected their Ellenberg scores to show limited relation to the drivers analysed. In total within our management categories there are 6665 measurements from 2717 plots surveyed for vegetation across 5 years and 588 squares, with 3797 measurements having co-located soil measurements (no soil samples were taken in 1990). 2383 of the measurements with both plant and soils data were categorised as low-intensity management.

**Figure 1:**
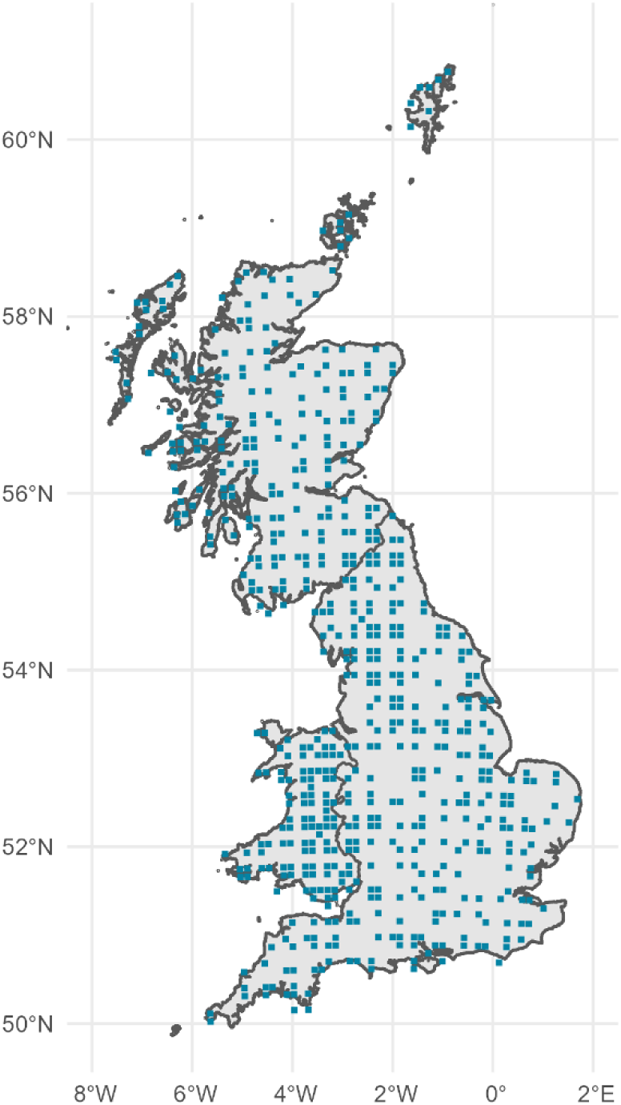
Map of survey square locations to the nearest 10km square.

Soil pH was measured in deionised water, and additionally in 2007 and 2019 soil pH was also measured in a CaCl_2_ solution. For a full description of the soils methods see Emmett et al (Emmett et al., 2008). All soil physicochemical data is available online at the NERC EIDC, see data availability statement. The measurement error of soil pH was calculated from randomly selected samples that were remeasured in 2007 and 2019. This procedure did not occur in 1978 and 1998, however due to a slight difference in the soil pH measurement procedure in 1998 there was a process of remeasuring samples in both 1998 and 2007 to compare the methods (Black et al., 2002; Emmett et al., 2010). The error from these remeasurements was used to create an expected error of the difference between 1978 and 1998, and 1998 and 2007 respectively.

Within every plot, Ellenberg R scores were calculated from the known species scores, with either the unweighted average or an average weighted by proportion of cover. This procedure was repeated for the full 14.14 by 14.14 m plots and the inner 2 by 2 m nest. An estimate of the error in these plant community variables was taken from the plots resurveyed as part of the Quality Assurance procedures in the survey years from 1990 onwards, the 1978 error is assumed to be the same as the 1990 error (see Supplementary Table 4 for all of the error estimates). In order to extract a standard deviation for use in the statistical modelling of the change in Ellenberg R the difference between the Ellenberg R score from the original and repeat survey was fitted to a normal distribution using the MASS package (Venables & Ripley, 2002). To compare to plant service provision the Ellenberg R scores of each plant species were compared to known values of service provision, including nectar-yielding species (Baude et al., 2016), agricultural forage grasses, crop wild relatives (Jarvis et al., 2015), and food plants for lowland birds and butterfly larvae (Smart et al., 2000). All data manipulation and statistical analysis was undertaken in R version 4.0.3 (R Core Team, 2020), plots were created using ggplot2 (Wickham, 2009).

### Atmospheric data sources

Atmospheric N and S deposition was taken from a 1km grid surface from the FRAME atmospheric chemistry transport model (Tipping et al., 2017; Tomlinson et al., 2021) and is available at the NERC EIDC, see data availability statement. Cumulative N deposition for the 8 years previous to survey was calculated for each square, as was the difference in S deposition over the 8 years previous to survey. Monthly rainfall data was taken from the HadUK-Grid 1km gridded climate observations provided by the Met Office (Met Office et al., 2020). Monthly rainfall over the month of sampling and the three months previous was averaged for every plot surveyed. The difference between the field season rainfalls was calculated for each year comparison.

### Statistical analysis

All statistical modelling presented here was undertaken in the brms package (v2.15.0) in R (v4.0.3), an interface to Stan (v2.21.2) (Bürkner, 2017; B. Carpenter et al., 2017). Ellenberg R and soil pH were both separately modelled as a response to an interaction between year (categorical) and the management intensity, with square identity as a group effect and a first order autoregressive process by year (sequentially numbered) and grouped by plot ID. The number of individual measurements in each model varied, with 6660 for the 4m^2^ unweighted Ellenberg R over 2717 plots, 6530 for the 4m^2^ cover weighted Ellenberg R over 2705 plots, and 6502 for the 200m^2^ Ellenberg R measurements over 2703 individual plots. For the soils data there were 3836 measurements of pH in water over 2323 plots, of which 2247 measurements were from the 21^st^ century and had corresponding pH in CaCl_2_ measurements. Pairwise contrasts between years were obtained using the emmeans package (v1.5.5-1, Lenth, 2021), and plotted using the tidybayes package (v2.3.1, Kay, 2020).

To investigate the relationship between changing Ellenberg R and soil pH, they were modelled in a multivariate Bayesian model with measurement error in the response variables being included. The change in Ellenberg R was modelled as a function of the change in soil pH, with an interaction with management intensity also included. The change in soil pH was modelled as a function of the change in sulphur deposition in the eight-year period before the second survey, the cumulative N deposition in the eight-year period before the second survey and the difference in pre-survey rainfall between the two time periods, to ensure standardisation of model coefficients all of these variables were scaled to have mean 0 and standard deviation of 1. The original means and standard deviations of these variables were −5.3 and 4.5 for S deposition, 134 and 72 for N deposition, and 8 and 30 for field season rainfall. All response variables were assumed to follow a Gaussian likelihood, and had square identity as a random effect, as well as an autoregressive component that accounted for time period where change occurred and had plot identity as a grouping factor. See supplementary methods for example code used to fit these models. We also explored the possibility of a non-linear relationship between S and N deposition and soil pH change but found it was not appropriate for our data, see Supplementary Methods for a discussion of this. Residuals from each model were plotted against broad habitat to check if there were systematic differences between habitat types (data not shown, no differences were found). The sample size varied per model due to differing numbers of small and large vegetation plots, being 1309 for the Ellenberg scores from the largest plots, and 1336 and 1458 for the weighted and unweighted Ellenberg scores from the 4m^2^ subplots respectively. Models were compared with 10-fold cross-validation using the loo package (v2.4.1, Vehtari et al., 2020).

## Results

### Trends in soil pH and Ellenberg R

Over the 40-year period surveys, soil pH increased in both high and low-intensity management habitats up until 2007, as reported previously from this data (Reynolds et al., 2013), but decreased in 2019 compared to 2007 (Figure 2a). Soil pH levels in 2019 were on average no different to the pH levels of 1978 and 1998 (Table 1) – note that no soils data was collected within the 1990 survey. However, the difference in acidity between 2007 and 2019 was much less apparent when pH was measured in a 0.01M CaCl_2_ solution, a method applied in the 21^st^ century surveys only. Those data provided no evidence for a drop in mean pH from 2007 to 2019 in low-intensity management habitats, and a smaller drop in pH in high-intensity habitats (Table 1, Supplementary Figure 1). Over the survey period plant communities under low-intensity management showed sustained increases in Ellenberg R, while high-intensity management land showed no overall trend in Ellenberg R (Figure 2b, Supplementary Figure 2). In order to evaluate the influence of the Ellenberg R calculation method on the trend estimate, Ellenberg R was calculated with and without cover weighting, and with or without constraining to species enumerated in the innermost nest of the plot. Measurement error was highest in the cover-weighted and smaller plot-based measurements. All four Ellenberg R metrics showed a continued increase in low intensity habitats, while remaining stable in high-intensity habitats (Supplementary Figure 2, Supplementary Table 1). This indicates the robustness of these trends to both measurement method and the variation in sample size, as more 4m^2^ plots were surveyed due to changes in survey protocol across the years. The measured changes in soil pH and Ellenberg R occurred during a period of dramatic decrease then stabilisation of S deposition (Figure 2c), and span a period during which N deposition initially increased before declining again (Figure 2d) but against a backdrop of at least 200 years of elevated N deposition (Fowler et al., 2005). Rainfall during the survey period also varied across the different years, with the highest levels of rainfall in 2007 and the lowest in 1990 (Supplementary Figure 3) In our dataset, higher Ellenberg R scores were associated with a greater diversity of plants and a higher number of species associated with multiple ecosystem functions, including nectar-yielding species, agricultural forage grasses, crop wild relatives, and food plants for lowland birds and butterfly larvae (Supplementary Figure 4).

**Figure 2:**
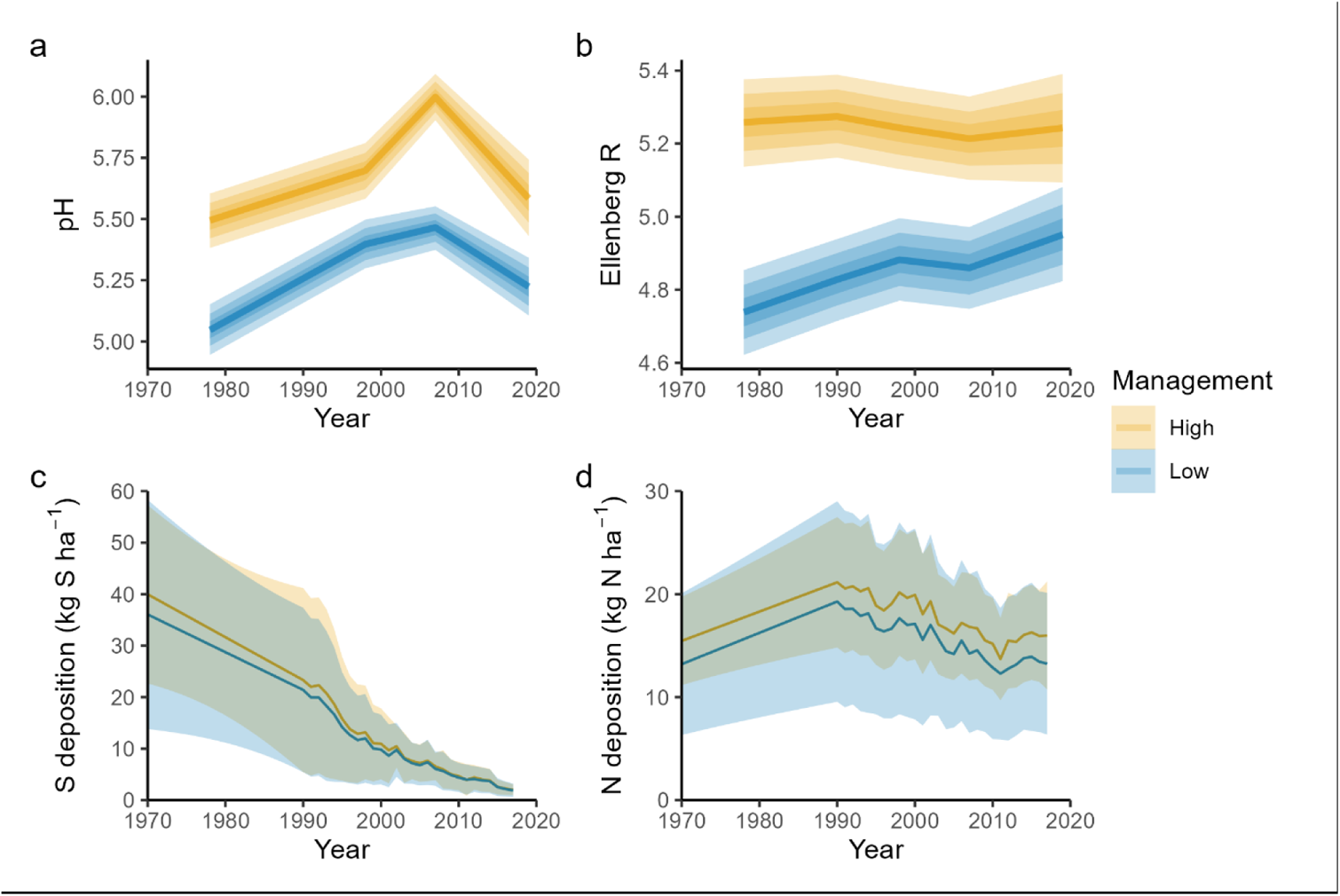
The change in soil pH and Ellenberg R (from the 4m^2^ plot and unweighted average) as modelled in response to management regime and year, represented by the median estimate and ribbons for the 50%, 80% and 95%tiles. Also shown are the changes in average S and N deposition across all of the plots in the two management categories, represented by a mean line and ribbons for the mean ± standard deviation. Note, 1km resolution S and N deposition estimates are not available between 1970 and 1990.

**Table 1:** Modelled difference between years for both pH in deionised water and pH in CaCl_2_ (most recent period only). The estimated difference and the lower and upper limits of the 95% highest posterior density (HPD) interval are given for each year contrast. Differences where the 95% HPD do not cross zero are in bold.

|  | Contrast | pH (DIW) |  |  | pH (CaCl<sub>2</sub>) |  |  |
| --- | --- | --- | --- | --- | --- | --- | --- |
|  |  | Estimate | Lower HPD | Upper HPD | Estimate | Lower HPD | Upper HPD |
| High intensity management | 1998-1978 | **0.203** | **0.110** | **0.294** | - | - | - |
|  | 2007-1978 | **0.504** | **0.417** | **0.598** | - | - | - |
|  | 2019-1978 | 0.092 | -0.063 | 0.231 | - | - | - |
|  | 2007-1998 | **0.302** | **0.206** | **0.388** | - | - | - |
|  | 2019-1998 | -0.112 | -0.262 | 0.037 | - | - | - |
|  | 2019-2007 | **-0.412** | **0.548** | **0.278** | **-0.141** | **-0.272** | **-0.007** |
| Low intensity management | 1998-1978 | **0.350** | **0.279** | **0.425** | - | - | - |
|  | 2007-1978 | **0.418** | **0.345** | **0.496** | - | - | - |
|  | 2019-1978 | **0.177** | **0.066** | **0.281** | - | - | - |
|  | 2007-1998 | **0.068** | **0.001** | **0.137** | - | - | - |
|  | 2019-1998 | **-0.173** | **-0.276** | **-0.065** | - | - | - |
|  | 2019-2007 | **-0.242** | **-0.337** | **-0.149** | 0.018 | -0.069 | 0.107 |

Overall, soil pH increased the most in the sites that showed the largest decreases in S deposition and the greatest cumulative N deposition (Figure 3). These two variables were negatively correlated such that the sites with the biggest decreases in S deposition also showed the largest cumulative N deposition (Pearson correlation −0.65). The estimated relationships between pH change and the two variables were broadly similar for the two management intensities. Changes in Ellenberg R were slightly negatively correlated with S deposition and slightly positively correlated with cumulative N deposition across all four Ellenberg R measurement methods, however the magnitude of this was small, with the greatest Pearson correlations being −0.10 and 0.04 for S and N deposition respectively with the unweighted score from the larger plot. For low intensity habitat only the magnitude was marginally higher, −0.14 and 0.08.

**Figure 3:**
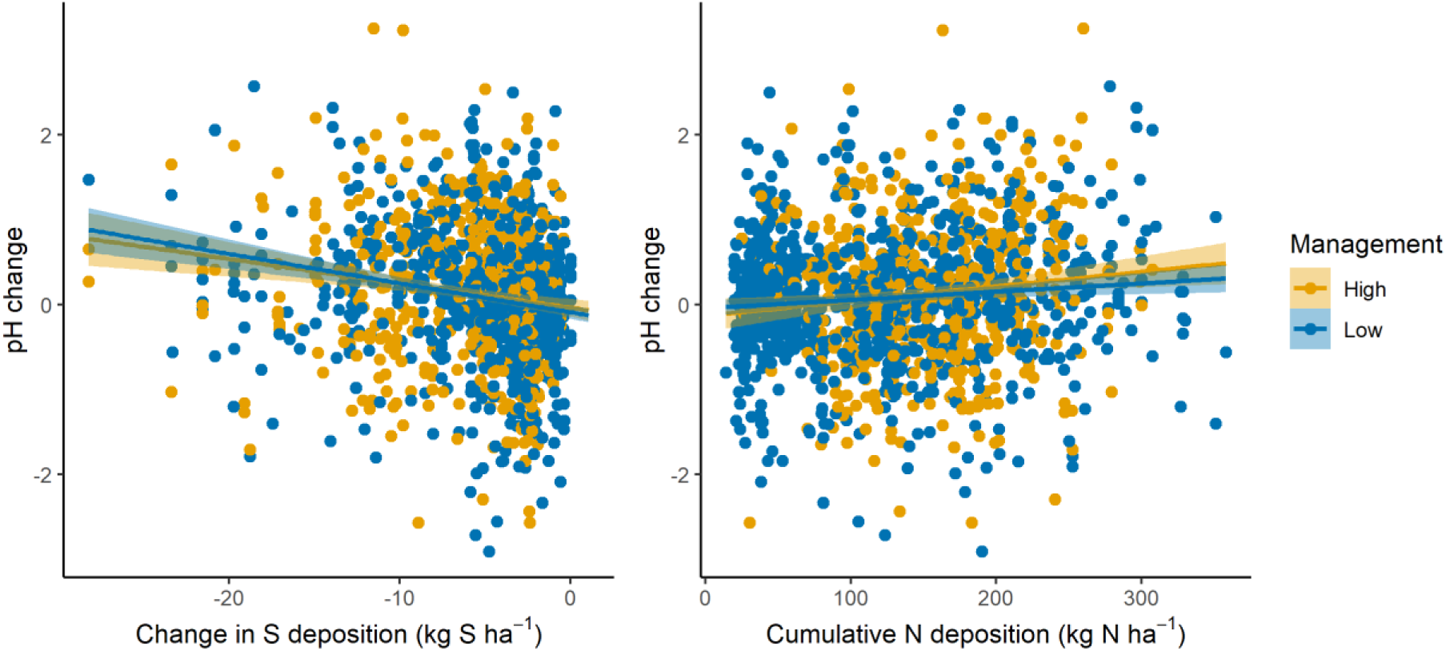
Change in pH between survey periods is negatively related to the change in S deposition (left) and positively related to cumulative N deposition (right). Both deposition statistics are from the 8 year period prior to the second survey. The points are the actual data, with each point representing a site/time period combination, and the lines plus ribbons are the estimated mean and the 2.5% and 97.5% quantiles of that mean.

### Soil pH change drives Ellenberg R change

Relationships between S and N deposition, soil pH and corresponding influences upon plant Ellenberg R scores were modelled in a multivariate Bayesian framework, where 1) change in S deposition, cumulative N deposition and difference in mean rainfall in the four months prior to and inclusive of survey were applied as predictors of the change in soil pH and 2) soil pH was used to predict the change in Ellenberg R. This enables us to evaluate the effect of changing atmospheric deposition upon soil pH while accounting for changes in precipitation. Both the soil pH and Ellenberg R models took into account differences in measurement error between the surveys, temporal autocorrelation at each site, and spatial autocorrelation within each 1km square. Temporal autocorrelation was estimated to be negative for both pH and Ellenberg R, indicating a “regression to the mean” effect. The hierarchical 1km square effect was greater in the soil pH model than in the Ellenberg R model, and there was no evidence of correlation between the square effects in the two models (Supplementary Table 2, Supplementary Figure 5).

We found a positive relationship between soil pH and Ellenberg R change, with an estimated increase in Ellenberg R of ~ 0.15 units per unit increase in soil pH (~30% of the interquartile range in Ellenberg R, Figure 4, Supplementary Figure 6, Supplementary Table 2). This was consistent across both management intensities and all four Ellenberg R metrics (Supplementary Figure 6). The weighted measurements of Ellenberg R appeared slightly more responsive to change in soil pH in the high intensity management habitats but this effect was small, while absent in the low intensity management habitat (Supplementary Figure 6). There was no effect of habitat type upon the model residuals, indicating no identifiable difference in the response of Ellenberg R to pH change, or pH change to atmospheric deposition, between woodland, grassland and heathland. If we had not taken into account measurement error in soil pH and Ellenberg R this relationship would have remained positive but at a smaller effect size of around 0.1 units of Ellenberg R change for a change in soil pH of 1 unit (Supplementary Table 3).

**Figure 4:**
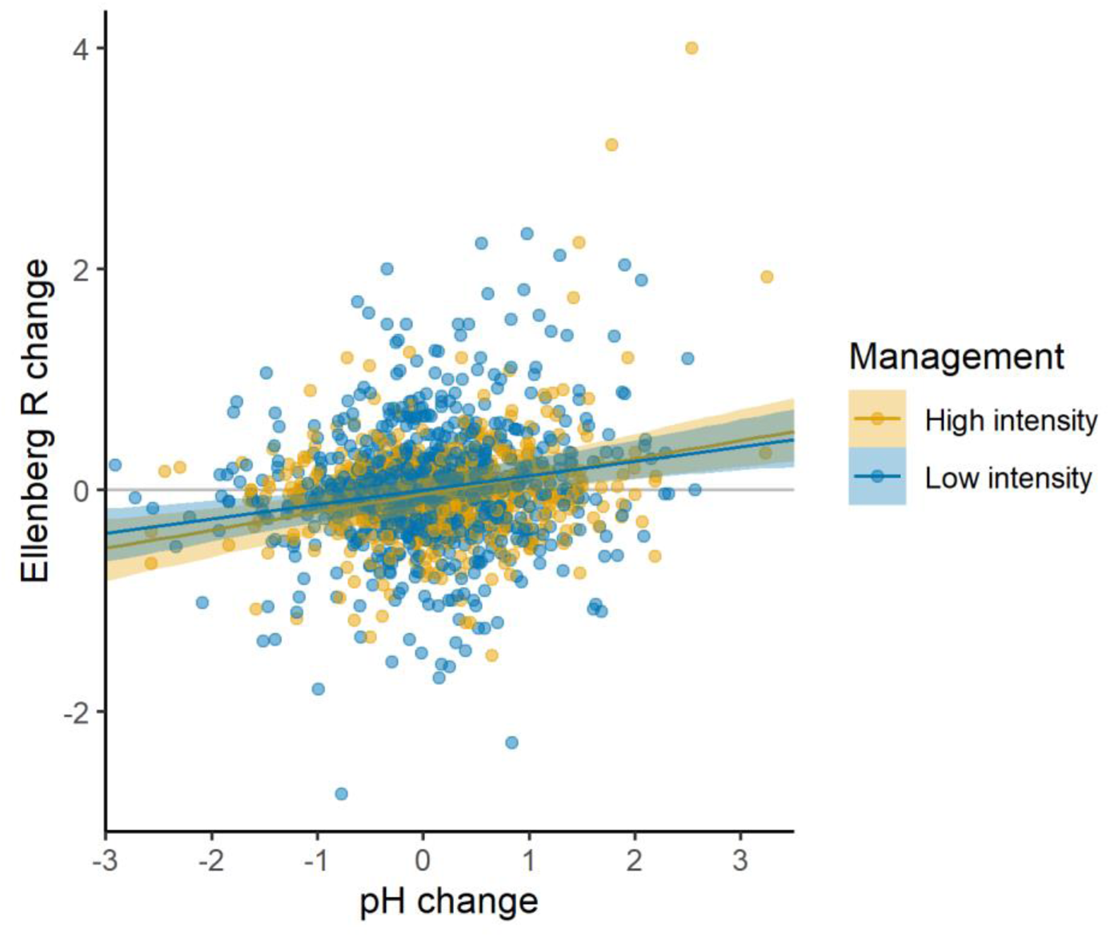
Soil pH change is positively related to Ellenberg R change within both high and low intensity management habitats. Data and model shown here are the Ellenberg R measured within the 4m^2^ plot and unweighted by cover. The points are the actual data and the lines plus ribbons are the estimated mean and the 2.5% and 97.5% quantiles of that mean.

Soil pH change was still slightly negatively associated with the change in S deposition in low intensity habitat once changes in field season rainfall and N deposition were accounted for. However, the effect was smaller for high intensity management areas and the 95% interval on the posterior parameter distribution included zero (Figure 5, Supplementary Figure 6). Soil pH change was positively associated with the difference in pre-survey rainfall between survey years in all four models, with an increase of around 0.1 pH units for every 30mm increase in rainfall across the field season (95% quantile 0.04-0.14, Supplementary Table 2). The N deposition effect was much reduced in magnitude once the S deposition effect and rainfall changes were accounted for (Figure 5, Supplementary Figure 6, Supplementary Table 2). The parameter estimates for the N and S deposition effects were positively correlated (Pearson correlation ~0.75 in all four models), meaning that when the model estimated a strong positive effect of N deposition this only occurred with a corresponding decrease in the estimated effect of S deposition. Adding direct linkages from N and S deposition to Ellenberg R resulted in no improvement in model out-of-sample prediction performance based on 10-fold cross validation (expected log-predictive density (ELPD) difference within one standard error of zero for all four models). The majority of the N and S parameter estimates were also very close to zero showing no particular residual effect of N and S deposition upon Ellenberg R change once pH change is accounted for (Supplementary Figure 7). However, within the unweighted Ellenberg R score for the whole 200m^2^ plot, in theory the most sensitive to change in rare species, there was some evidence for a direct negative effect of S deposition on Ellenberg R change in low intensity management habitats only (Supplementary Figure 7).

**Figure 5:**
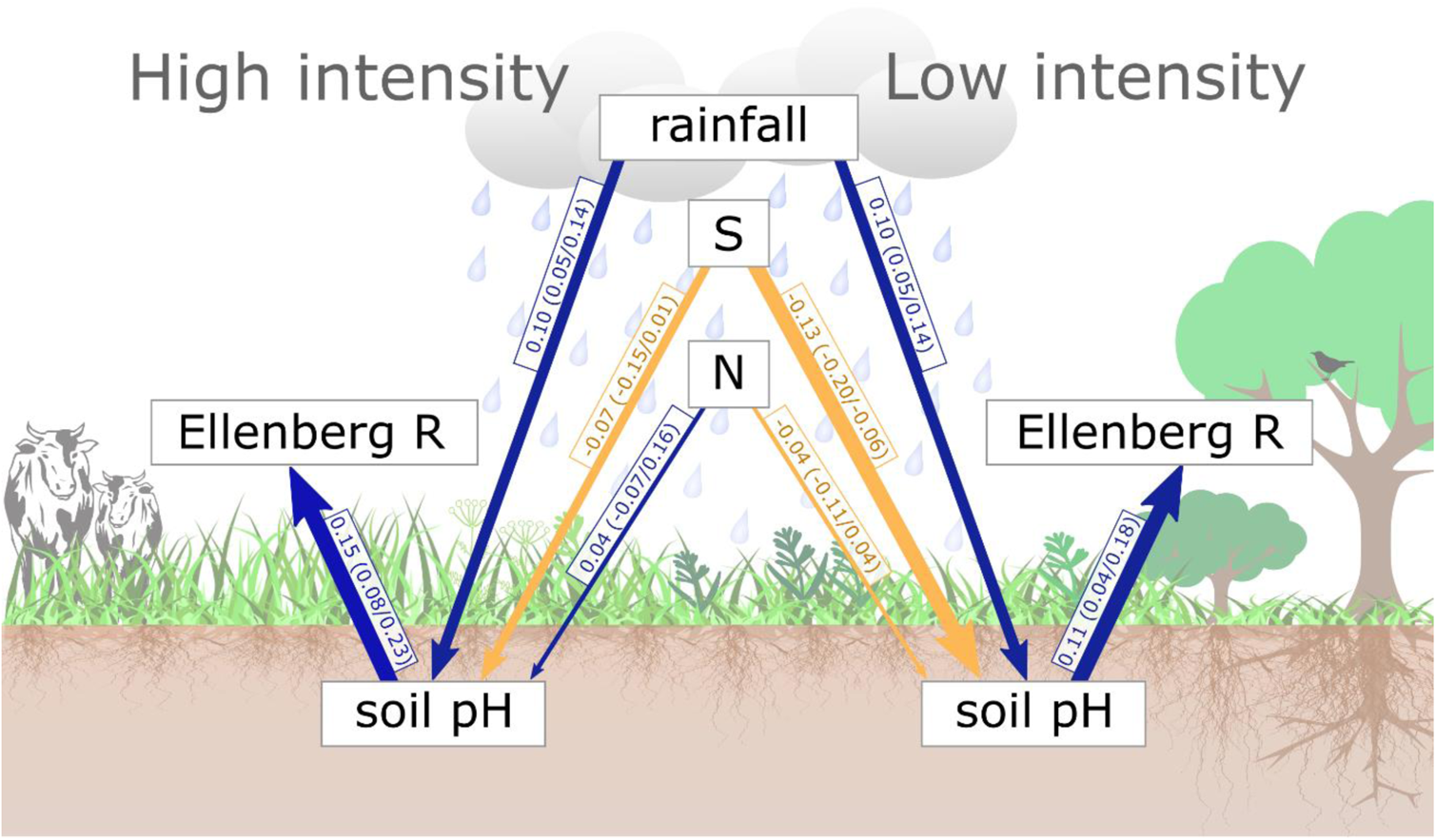
Graphical representation of the multivariate model results (Ellenberg R score from the unweighted 4m^2^ plot) with results from high intensity management land on the left and low intensity management land on the right, where each arrow represents a specific model parameter and is labelled with the median parameter estimate (2.5%/97.5% quantiles). The rainfall effect was assumed to be constant in the two habitat categories but is shown separately for graphical clarity. Blue arrows represent positive median relationships and yellow negative, note that the 95% interval contains zero for the N deposition arrows and the S deposition arrows in high intensity habitats.

## Discussion

Plant communities in semi-natural habitats have shown progressive recovery from acidification over the past 40 years, demonstrating both the promising reality of recovery from pollution impacts within semi-natural ecosystems as well as the longer time scales required to see the positive ecological impacts of legislative change. Our results clearly demonstrate that air pollution control policy can lead to ecological recovery even though reductions in S deposition may have been initiated by economic, rather than ecological, considerations (RoTAP, 2012). As expected, the pattern of recovery from acidification was not apparent in the more highly managed pasture habitats which showed relatively stable acidity preference over the past 40 years. This indicates that in both the above and below-ground parts of the ecosystem, intensive land management effects and higher pH starting conditions have to an extent eclipsed atmospheric deposition effects. However, this does not mean that these habitats have not responded to changes in soil acidity over time as we still found a clear link between soil pH change and Ellenberg R change in intensive grasslands. It appears that the lack of evidence for a direct link between plant community response and decreasing S deposition in intensive pasture to is likely due to soil pH changes not being as strongly linked to S deposition changes in these systems.

Increases in soil pH across GB over the late 20^th^ and first decade of the 21^st^ century appear to have stalled, or in some case reversed, in the past decade. We can be confident of the decreasing pH in high-intensity management areas which was found in both soil pH in water and in CaCl_2_, however low-intensity management habitats showed no change in soil pH in CaCl_2_ versus a decline in soil pH in water. Our data can be compared to a previous study of soil pH from farm survey data which indicated that soil pH has recently decreased in the north-west of England and Wales (Rawlins et al., 2017). The increasing acidity in land managed for pasture is potentially related to the decline in liming of agricultural soils across the UK, which is now below the level required to maintain soil pH in the productive zone (Goulding, 2016). Increasing liming or alternatively applying basalt to crop and pasture land could therefore increase pH again, and in the latter option also provide a valuable service in carbon capture (Beerling et al., 2018). However, liming of agricultural soils in the UK dropped in the 1990s and has remained at a fairly constant level since 2000 (Holland et al., 2018; The British Survey of Fertiliser Practice, 2020), indicating that there may be other factors involved in the difference between 2007 and 2019 soil pH such as changes in fertiliser use. There is also the possibility that climate change may be impacting the recovery from acid deposition, as both changing temperature and precipitation have been found to impact soil pH and interact with S deposition, moreover, we also found an effect of rainfall upon pH change (Adamson et al., 2001; Chen et al., 2021; Li et al., 2019). Rainfall could be affecting pH through increasing soil moisture leading to a shift from the dominance of nitrification processes under aerobic conditions towards increasing anaerobic conditions and thus increasing proton consumption processes which promote N_2_O emissions and increasing pH (Davidson et al., 2000; Dobbie & Smith, 2001; Russow et al., 2000). However, the extent of the difference between the predicted pH changes from the two different measurement methods–in water versus in CaCl_2_–raises the issue of whether our measured pH change optimally represents changes in “plant-visible” soil acidity. Soil pH measurements made in water are known to be more affected by salt solution and therefore soil moisture effects (Hester & Shelton, 1933; Kissel et al., 2009). Measurement of soil pH in CaCl_2_ should be more robust to changes in soil moisture, and more closely relates to the ionic strength of the near-root and near-microbe surfaces, and so would be more suited for tracking changes over time in soil acidity relevant to plant communities and soil functions (Custos et al., 2020; Schofield & Taylor, 1955). Unfortunately we do not have pH in CaCl_2_ measurements for the entire soil series as this was first measured in 2007. Instead we must consider carefully the role of soil moisture in driving the patterns of increasing pH in the early part of the time series. While field season rainfall across the UK in 1978, 1998 and 2019 field seasons was broadly similar, the 2007 season was on average wetter which may explain some of the increase in soil pH. Integrating rainfall measurements into our multivariate modelling allowed us to demonstrate how both S deposition and rainfall together influenced the soil pH measured over these time periods illustrating individual and additive roles for each driver. This also implies that ongoing impacts of reduced atmospheric deposition are likely to be modified by a backdrop of ongoing changes in precipitation again emphasising the importance of accounting for interactions between global change drivers when attributing signals of ecosystem change (Perring et al., 2018).

Our analysis has shown that the major drivers of soil pH change over the past forty years are decreasing S deposition and variation in rainfall. Rainfall was found to be strongly positively related to soil pH change such that a wetter field season had higher soil pH on average, consistent with our knowledge of soil chemistry and previous studies on the effect of seasonality upon soil pH (Hester & Shelton, 1933; Kissel et al., 2009; Marwanto et al., 2018). Inclusion of this rainfall term also enables us to have greater confidence that the S deposition effect we have found is representative of a direct effect once changes in climate and therefore soil moisture are accounted for. While we found a strong effect of the decreases in S deposition upon soil pH, we found no clear evidence of an additional eutrophying effect of cumulative N deposition on plant species composition where soil pH had increased. It is possible that the eutrophication effect on Ellenberg R may be only slightly positive or non-linear and thus not detected by our analysis, however the noisiness of our data and the difficulties we encountered in fitting non-linear models (see Supplementary Methods) indicate that this dataset would not be able to distinguish the subtle non-linear relationships implied by this hypothesis from ecological noise.

Our multivariate modelling demonstrates the close linkages between soil pH and plant species compositional change, tracked by the Ellenberg R indicator, with atmospheric deposition effects on the plant communities mediated by the changes in soil condition. The potential exception to this full mediation was the direct responsiveness of Ellenberg R to S deposition when the R score is measured as an unweighted average of the larger plot size. This particular formulation of the Ellenberg R score is more sensitive to rarer species and is also likely to be the least spatially coupled to the single point measurement of soil pH as it gives equal weight to plant species that are found in small numbers far from the soil sampling location. As soil pH experiences high levels of spatial variation at a local scale (Ball & Williams, 1968), it may be that this link between S deposition and Ellenberg R change appears to be separate from soil pH change only due to this decoupling of the Ellenberg R and soil measurements. Surprisingly, we found only a limited interaction between the management intensity and the Ellenberg R ~ pH relationship, despite having found divergent trends in soil pH and Ellenberg R change over time in high-intensity management habitats. The stability of Ellenberg R from 2007 to 2019 while soil pH decreased in high-intensity habitats could be related to lag effects or indicative of less close coupling of soil and vegetation change under intense management. However there was no indication of less close coupling of soil and vegetation change in the multivariate model results which showed a continued relationship between soil pH and Ellenberg R change in high intensity habitats. This was also unexpected due to the curvilinear relationship between Ellenberg R and soil pH, such that above pH 6 there tends to be a flattening of the Ellenberg R relationship with soil pH (Diekmann, 2003). Potential explanations for the persistence of the relationship between soil pH and Ellenberg R change in high-intensity management habitats include the co-occurrence of soil pH and plant community interventions in highly managed areas, such that as the soil is limed there is also introduction of high Ellenberg R plant species within seed mixes. It should also be noted that there was a lower response of soil pH change to S deposition within these high-intensity management habitats, indicating that acidity changes in both soil pH and plant communities are less affected by atmospheric deposition. Therefore, despite there being no overall trend in Ellenberg R in high-intensity pasture over the time period studied we can still conclude that the high-intensity pasture sites that did undergo a change in Ellenberg R were on average sites that showed equivalent change in soil pH.

While we identified a clear response by Ellenberg R to soil acidity, as originally intended by Ellenberg’s scheme, species’ Ellenberg R scores also tend to correlate with Ellenberg N (fertility) scores in a non-linear fashion (Diekmann, 2003; Diekmann & Dupré, 1997). Therefore, as the plants that prefer acidic environments also prefer low-fertility environments, the changing Ellenberg R may be reflecting the dual drivers of change of decreasing acid rain and increasing fertilisation (Maskell et al., 2010; Peppler-Lisbach et al., 2020). We attempted to explore this possibility by inclusion of N deposition within our modelling framework, which represented changes in atmospheric fertilisation of plant communities. Previously it has been proposed that decreasing acidity in aquatic systems may unlock a eutrophication effect such that the impacts of N deposition are greater as S deposition decreases (Schneider et al., 2018). This might also be expected to occur in land ecosystems as the impacts of aluminium toxicity decrease once pH is raised and macronutrient availability maximised above 5-6 (Jones et al., 2019; Zhao & Shen, 2018). However, we found no evidence of effects of N deposition upon either soil pH or directly upon Ellenberg R change. While the range of our estimated N deposition is large enough to include some possibility of an effect, on balance the likelihood is that Ellenberg R change is driven by soil acidity change rather than changes in N fertilisation. This could partly be due to the measured N deposition in the 8 years prior to the survey not being representative of either historical N deposition, or the N accumulated within the soil. It is possible that phosphorus fertilisation or limitation could be linked to changes in Ellenberg R as phosphorus is known to differentially influence plant species of varying Ellenberg N scores (Löfgren et al., 2020). Use of phosphorus fertilisers has declined in the UK and there is some evidence of a decline in available phosphorus in Scottish soils from the 1980s to the present, which would affect both the fields directly fertilised and also leaching to the surrounding landscape (Lilly et al., 2020), however analysis of changes from the 1990s to the present within data collected from large-scale surveys and to support farm management has not found a change in available phosphorus in UK soils (Edwards et al., 2016; Rawlins et al., 2017; Seaton et al., 2021). Residual variation in Ellenberg R change over these decades could be explained by the local-scale changes in nutrient availability, biotic interactions, and land use that we were unable to consider in this broad-scale analysis.

The inclusion of measurement error within our modelling framework allowed for identification of a stronger relationship between soil pH change and Ellenberg R change than would have been found otherwise. Measurement error affects every part of the sampling to analysis pipeline and it has frequently been found to be extensive and pervasive in ecological and soil science (Desaules, 2012; Morrison, 2016). Within our data there were greater levels of measurement error within the Ellenberg R estimates that were weighted by plant cover, and in the smaller plots. Taking all of these errors into account resulted in robust relationships between Ellenberg R change both over time and in relation to pH change across all four measurement methods of the plant community. While we might expect the Ellenberg R scores weighted by plant cover to be more responsive to environmental change it may be that this is balanced out by the increased error of these measurements (Diekmann, 2003), which could explain previous results showing no greater correlation of cover weighted Ellenberg scores with environmental variables (Carpenter & Goodenough, 2014; Käfer & Witte, 2004). Inclusion of measurement error within our analysis did limit the analysis we could perform due to the limited quality assurance data from the early survey years, however we consider this to be well worth it for the increased realism of our results.

Our results indicate that positive effects on plant biodiversity and ecosystem service provision are likely to have resulted from the causal chain linking reduced S deposition to change in our index of plant acidity preferences. As this mean acidity index increases, conditions becomes more favourable to a larger pool of species. This also results in passive selection for larger subsets of species valued because of the support they provide to ecosystem services and functions (Supplementary Figure 4). These groups include plants that underpin positive services such as crop wild relatives (Jarvis et al., 2015), nectar-yielding species (Baude et al., 2016), agricultural forage grasses and food plants for lowland birds and butterfly larvae (Smart et al., 2000). However, there are other plant groups less beneficial to human activities such as injurious weeds that are also more common at higher Ellenberg R scores (Maskell et al., 2020). Overall, these changes in plant service provision indicate how changing atmospheric deposition not only influences soil acidity and thus plant composition but also acts through these changes in plant communities to influence the health of the whole ecosystem, including bird and pollinator communities.

The close linkages between the plant community and the soil chemical status we have found here demonstrate the range of potential future impacts of a variety of environmental stressors. Soil pH is an emergent property of the soil system that is responsive to a variety of factors as well as influencing a wide range of both microbially- and chemically-driven soil functions (Neina, 2019). The range of environmental stressors that ecosystems are currently experiencing can affect some elements of the soil and plant system more than others, e.g. soil fungal communities are known to be particularly responsive to synergistic effects of multiple stressors (Rillig et al., 2019). These changes in microbial communities could then feed back to soil pH changes with knock on effects and positive feedback loops influencing soil and plant health. Moreover, it is clear that changes in soil pH driven by anthropogenic changes in atmospheric deposition and land management will have concomitant impact on soil and general ecosystem health. Of particular concern is the potential for changes to occur on different timeframes, as seen here within the changes in the plant community over time relative to changes in soil pH. There have also been observed changes in the soil pH ~ carbon relationship over time in GB soils (Thomas et al., 2020). This raises the possibility of a change in the mechanisms that drive soil and ecosystem functions and increasing disconnect between differing ecosystem components, such that future change will become more difficult to predict and mitigate (Hawkes & Keitt, 2015; M. D. Smith et al., 2009).

## Acknowledgements

This work was funded by the Natural Environment Research Council (part of UK Research and Innovation) under the UK-SCAPE Programme delivering National Capability (Grant Reference NE/R016429/1). Countryside Survey was funded by a consortium of funders led by the Natural Environment Research Council and the Department for the Environment Food and Rural Affairs, full list available at the Countryside Survey web-page (countrysidesurvey.org.uk).

## Author contributions

SMS conceived this project, with FMS and SMS designing the analyses with input from DAR, DM, IL and BAE. DAR and IL provided the soils data, SMS the plant data and ST the atmospheric deposition data. FMS carried out the statistical analysis with assistance from PB. FMS wrote the first draft of the manuscript, with subsequent revisions by SMS, DM, DAR, ST, PB, and IL. All authors read and approved the final draft of this manuscript.

## Materials and correspondence

All correspondence and requests for materials should be addressed to Fiona Seaton.

## Data availability statement

All vegetation, soils and atmospheric deposition data is available for download from the NERC Environmental Information Data Centre, with digital object identifier links detailed below for each dataset. The error estimates for the Ellenberg R and soil pH change over time are included in Supplementary Table 4.

### Vegetation data

Barr, C. J. et al. Countryside Survey 1978 vegetation plot data. (2014) doi: 10.5285/67bbfabb-d981-4ced-b7e7-225205de9c96.

Barr, C. J. et al. Countryside Survey 1990 vegetation plot data. (2014) doi: 10.5285/26e79792-5ffc-4116-9ac7-72193dd7f191.

Barr, C. J. et al. Countryside Survey 1998 vegetation plot data. (2014) doi: 10.5285/07896bb2-7078-468c-b56d-fb8b41d47065.

Bunce, R. G. H. et al. Countryside Survey 2007 vegetation plot data. (2014) doi: 10.5285/57f97915-8ff1-473b-8c77-2564cbd747bc.

Smart, S. M. et al. Vegetation plot data from the UKCEH Countryside Survey, Great Britain, 2019. (2020) doi: 10.5285/fd6ae272-aeb5-4573-8e8a-7ccfae64f506.

### Soils data

Bunce, R. G. et al. Soil physico-chemical properties 1978 [Countryside Survey]. (2016) doi: 10.5285/85c71959-0f7c-4f04-b3a7-152673107a85.

Black, H. I. J. et al. Soil physico-chemical properties 1998 [Countryside Survey]. (2016) doi: 10.5285/9d1eada2-3f8b-4a7b-a9b0-a7a04d05ff72.

Emmett, B. A. et al. Soil physico-chemical properties 2007 [Countryside Survey]. (2016) doi: 10.5285/79669141-cde5-49f0-b24d-f3c6a1a52db8.

Robinson, D. A. et al. Topsoil physico-chemical properties from the UKCEH Countryside Survey, Great Britain, 2019. (2020) doi: 10.5285/aaada12b-0af0-44ba-8ffc-5e07f410f435.

### Atmospheric deposition data

Tomlinson, S. J. et al. Nitrogen deposition in the UK at 1km resolution, 1990-2017. (2020) doi: 10.5285/9b203324-6b37-4e91-b028-e073b197fb9f.

## Supplementary

### Supplementary methods

We wished to test the assumption of linearity inherent in our modelling so used penalised thin-plate splines to represent the impact of S and N deposition upon soil pH (Wood, 2003), fit as separate splines for each management category. We found evidence of funnel relationships between the log standard deviation and the location parameters of the penalised splines, a particular complex posterior geometry (Betancourt, 2017), which led to many divergent transitions within the Hamiltonian Monte Carlo (HMC) algorithm implemented in Stan. This was particularly evident when splines were fit to S deposition without any interaction with other parameters. Reducing the stepsize of the HMC algorithm through increasing the adapt_delta hyper-parameter to 0.99999999 removed the divergent transitions. Examination of the conditional smooths plot of the deposition effects upon soil pH showed largely linear responses (Figure A below). These largely linear relationships and the persistent funnels motivated our decision to model the responses of pH and Ellenberg R to S and N deposition as linear models.

**Figure A:**
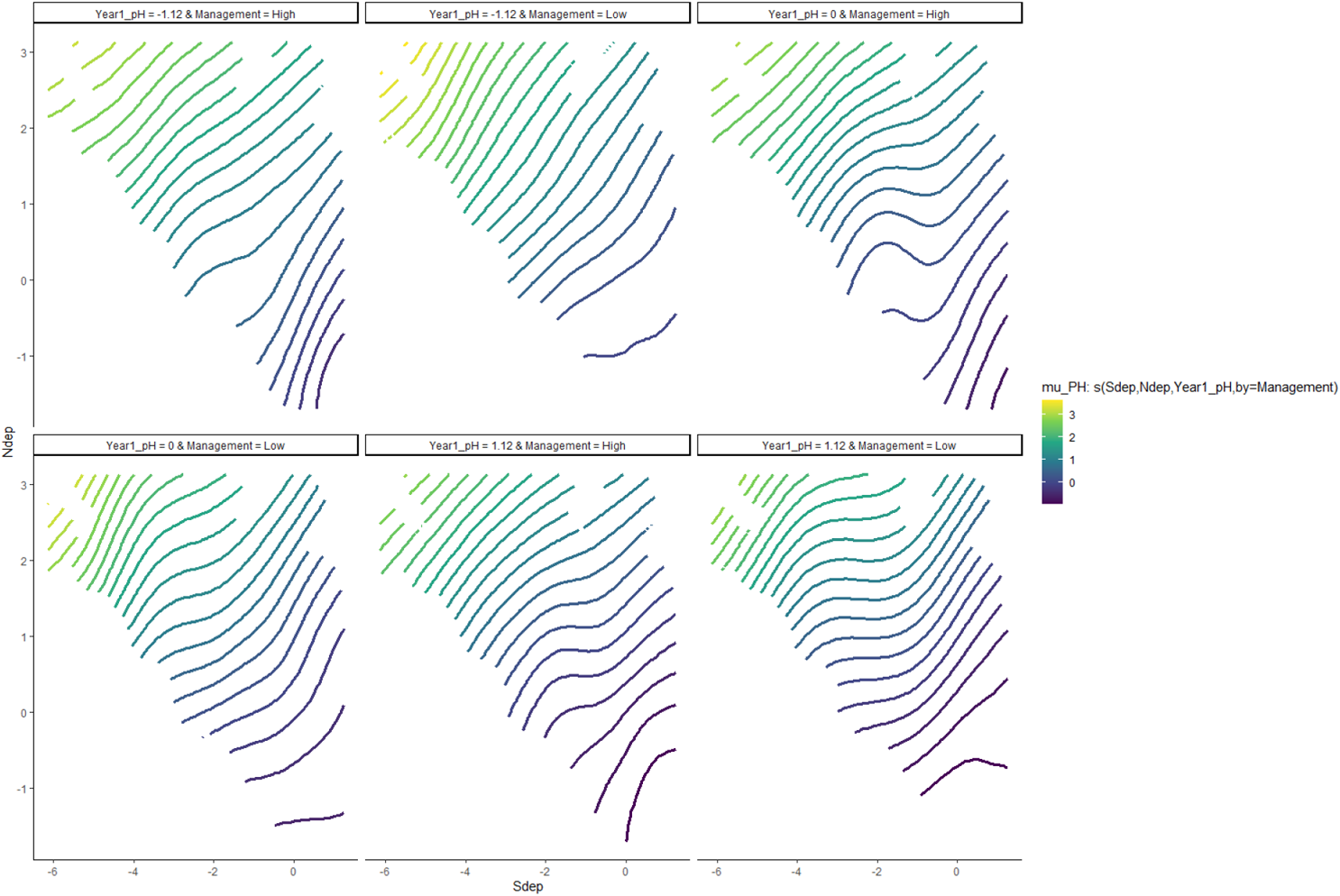
Contour plots of the splines showing the largely linear response (parallel contour lines) of pH to a spline upon S deposition, N deposition for differing initial pH values and management values.

### Model code

The following section gives the priors and model code used to fit the models within brms (version 2.15.0, rstan version 2.21.2).

Year models without measurement error:

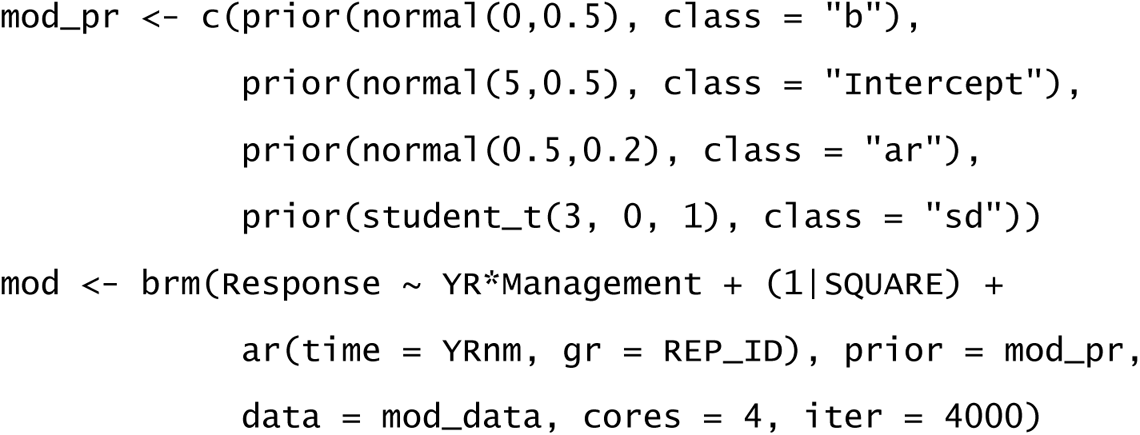

Year models with measurement error:

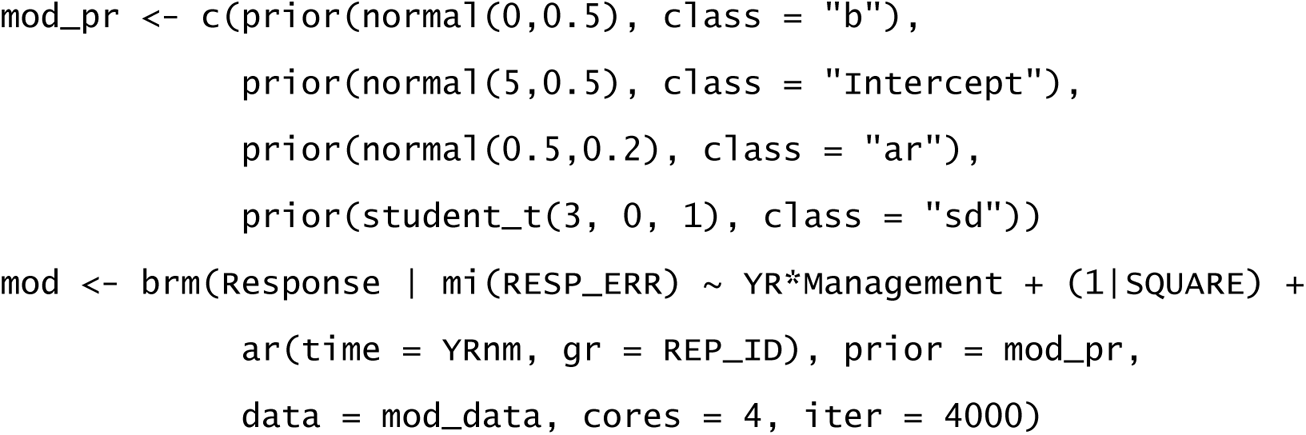

pH ~ Sdep OR Ndep models:

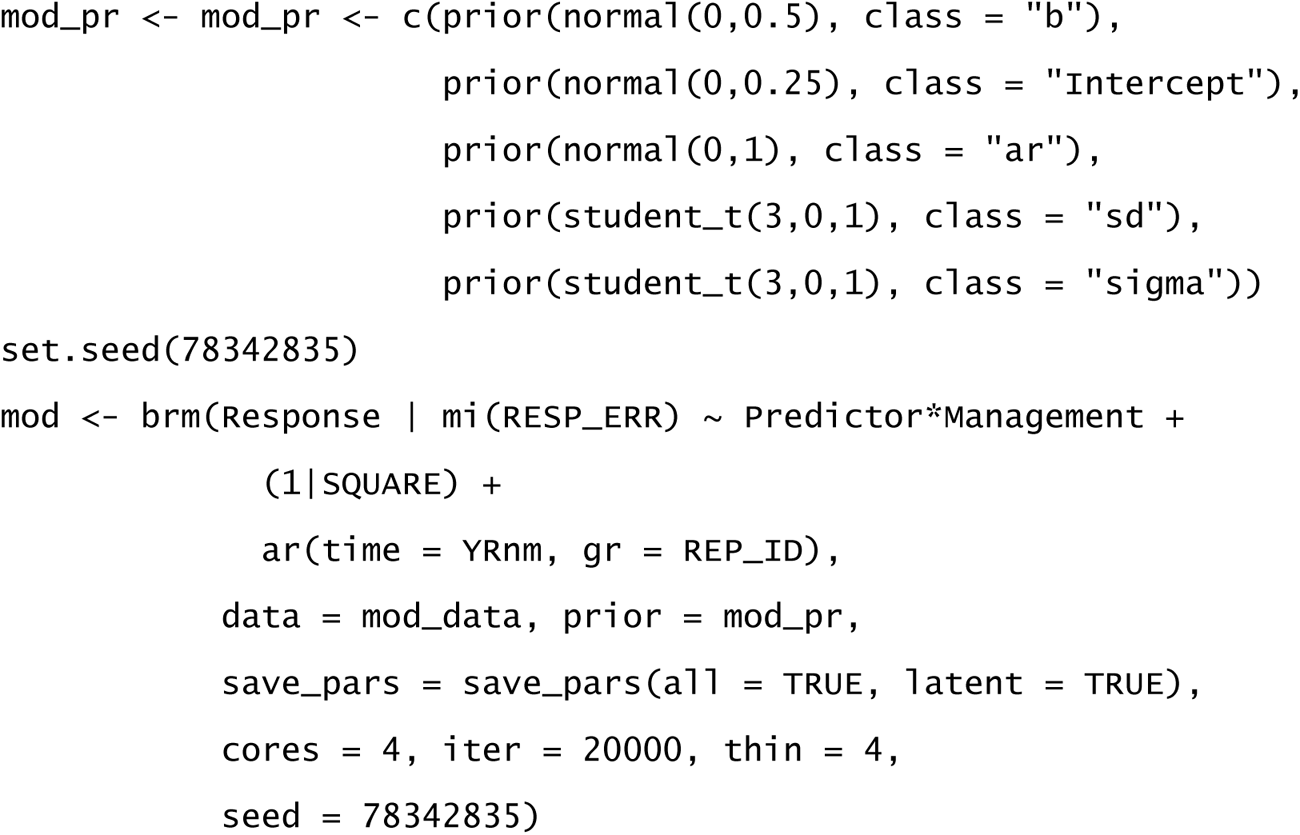

Multivariate model:

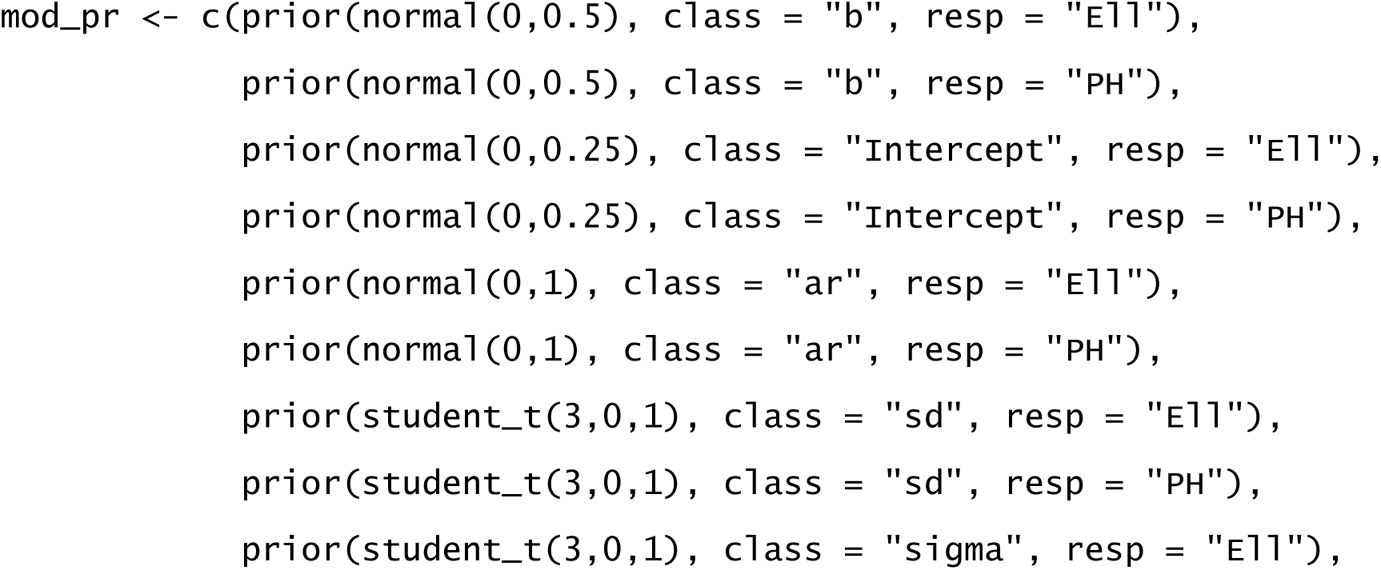

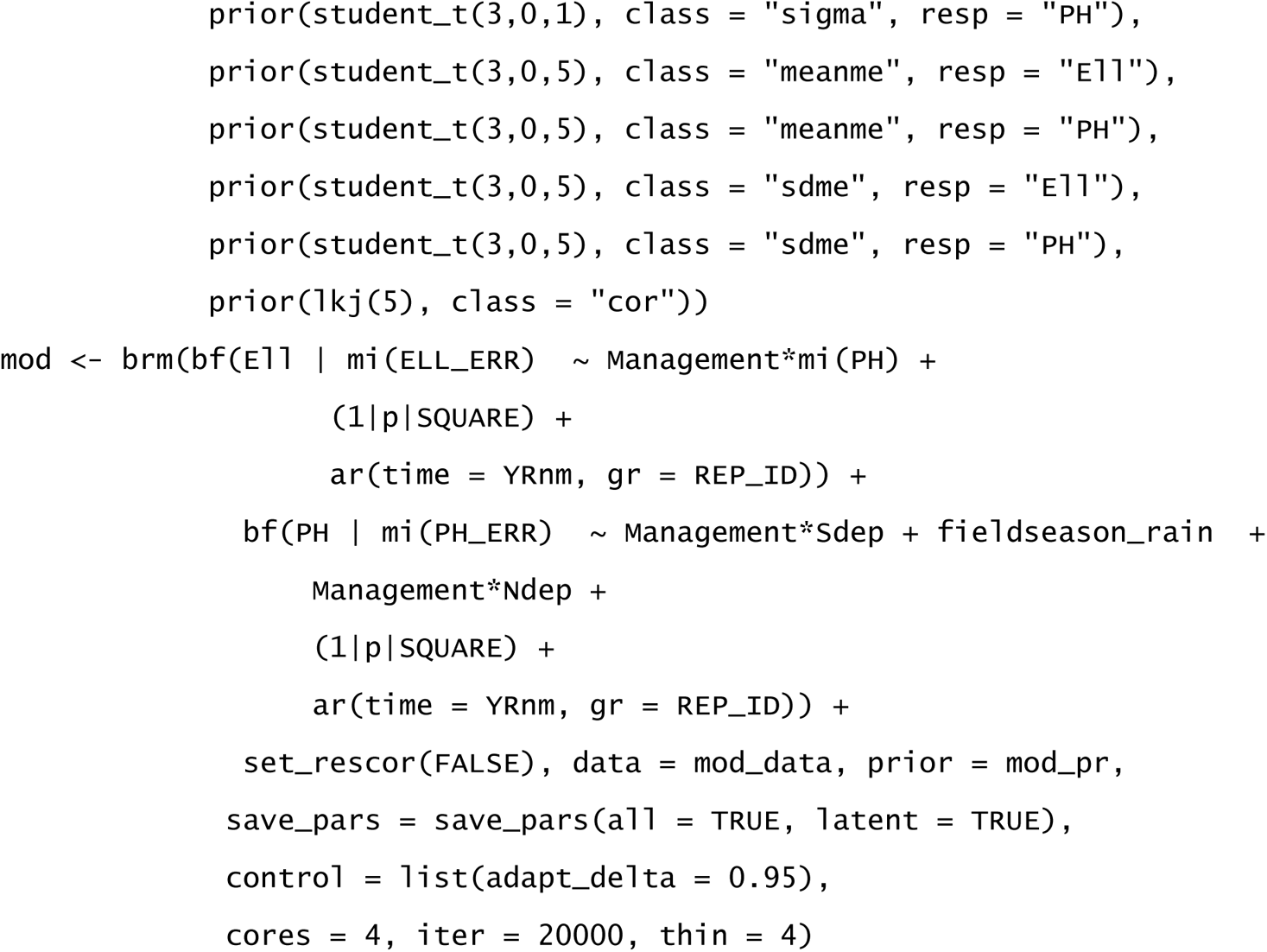

Three multivariate models (out of eight) had a small number of divergent transitions, however no pathological model structures were seen when examining the parameter plots and therefore we increased adapt_delta to 0.95 which removed all of these divergent transitions. We also ran versions of the multivariate models with N and S deposition as direct predictors of Ellenberg R change – each with an interaction with management – and versions of all of these models without measurement error included.

## Supplementary Figures

**Figure 1:**
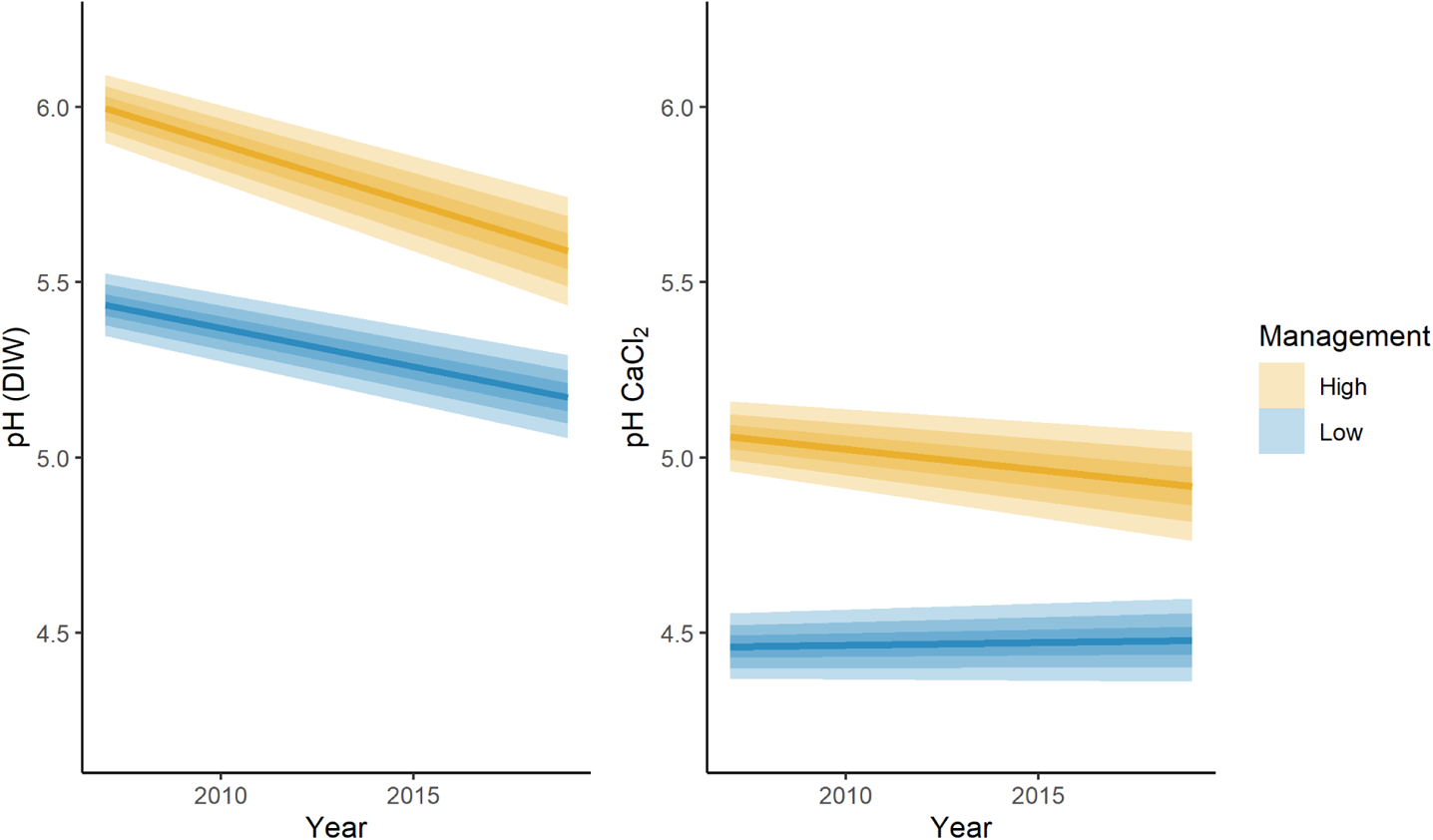
The change in soil pH from 2007 to 2019 when measured with deionised water vs calcium chloride solution.

**Figure 2:**
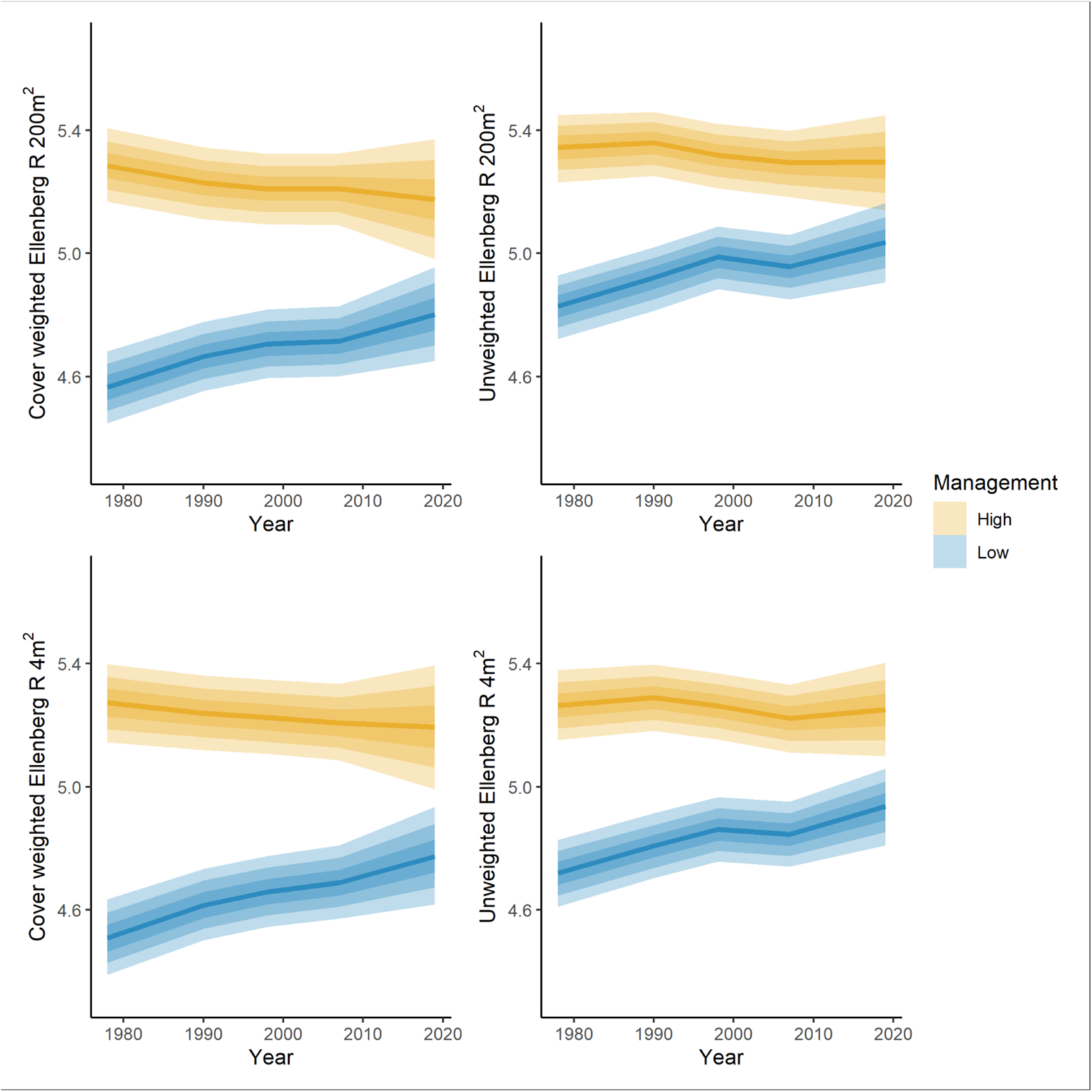
Predicted mean Ellenberg R scores for the two different management categories per year of survey for each Ellenberg measurement method. Results from the model including measurement error

**Figure 3:**
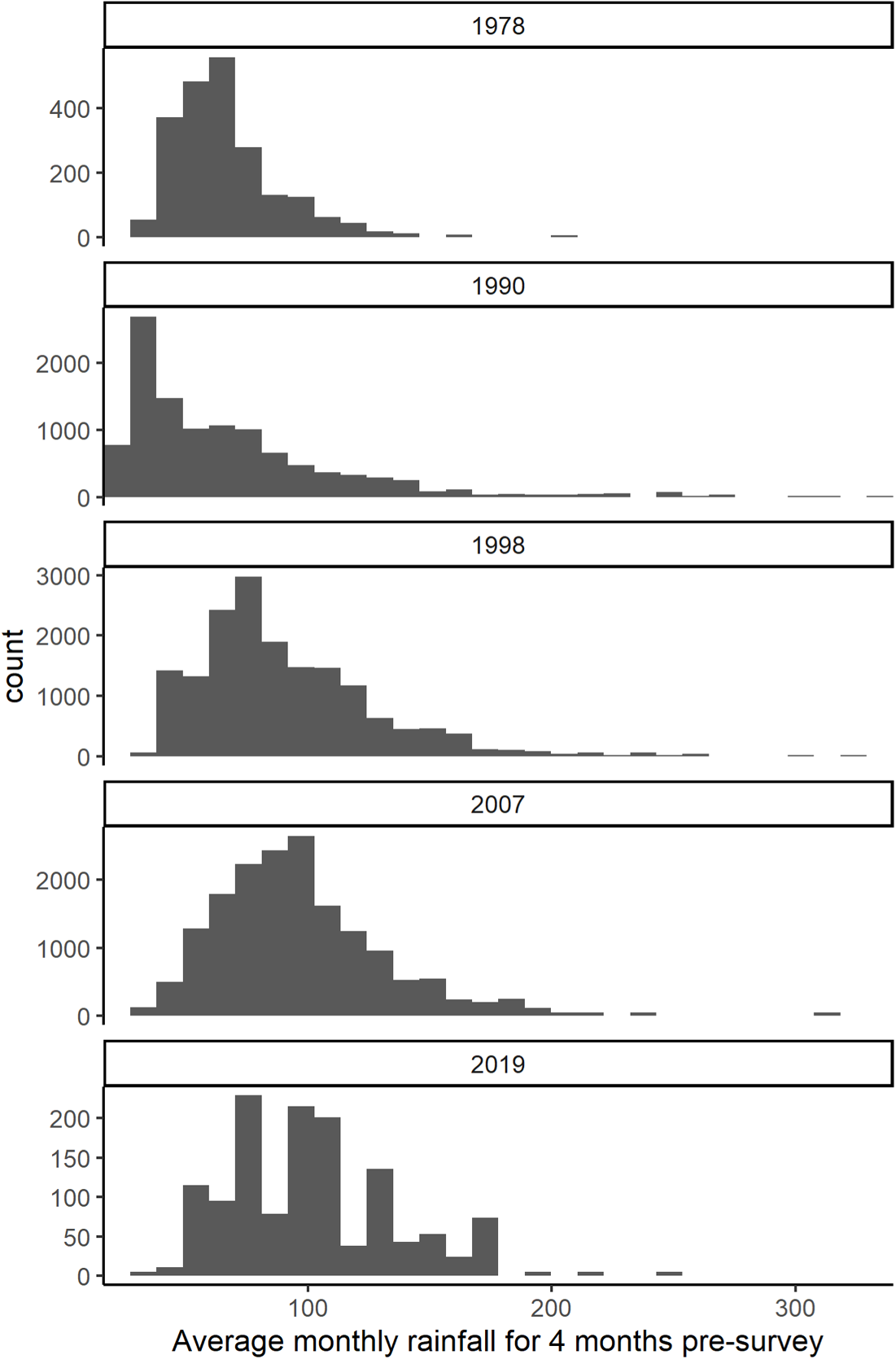
The variation in average monthly rainfall for the 4 months pre-survey between the different survey years.

**Figure 4:**
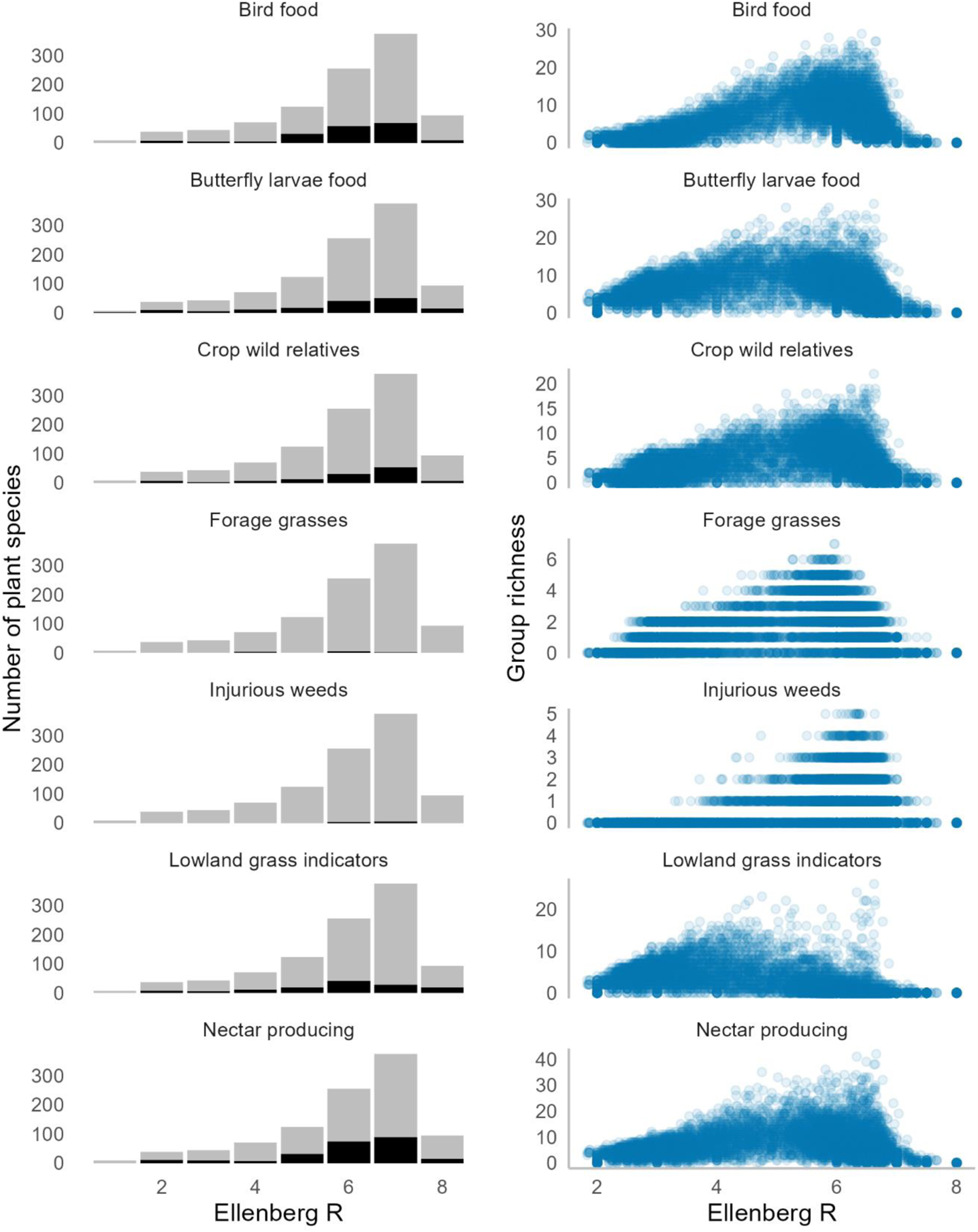
The proportion of plant species that carry out some function within each Ellenberg R category (left) and the richness of the functional group plotted against Ellenberg R score for all sites (right). The black bars show the species that do carry out the function, which are compared to the light grey bars which show the number of species that do not carry out the function. Functional plant categories are in order from the top: plant species that are food plants for lowland birds (Smart et al., 2000), food plants for butterfly larvae (Smart et al., 2000), crop wild relatives (Jarvis et al., 2015), forage grass species, injurious weeds, Common Standards Monitoring lowland grass quality indicators, and nectar producing plants (Baude et al., 2016).

**Figure 5:**
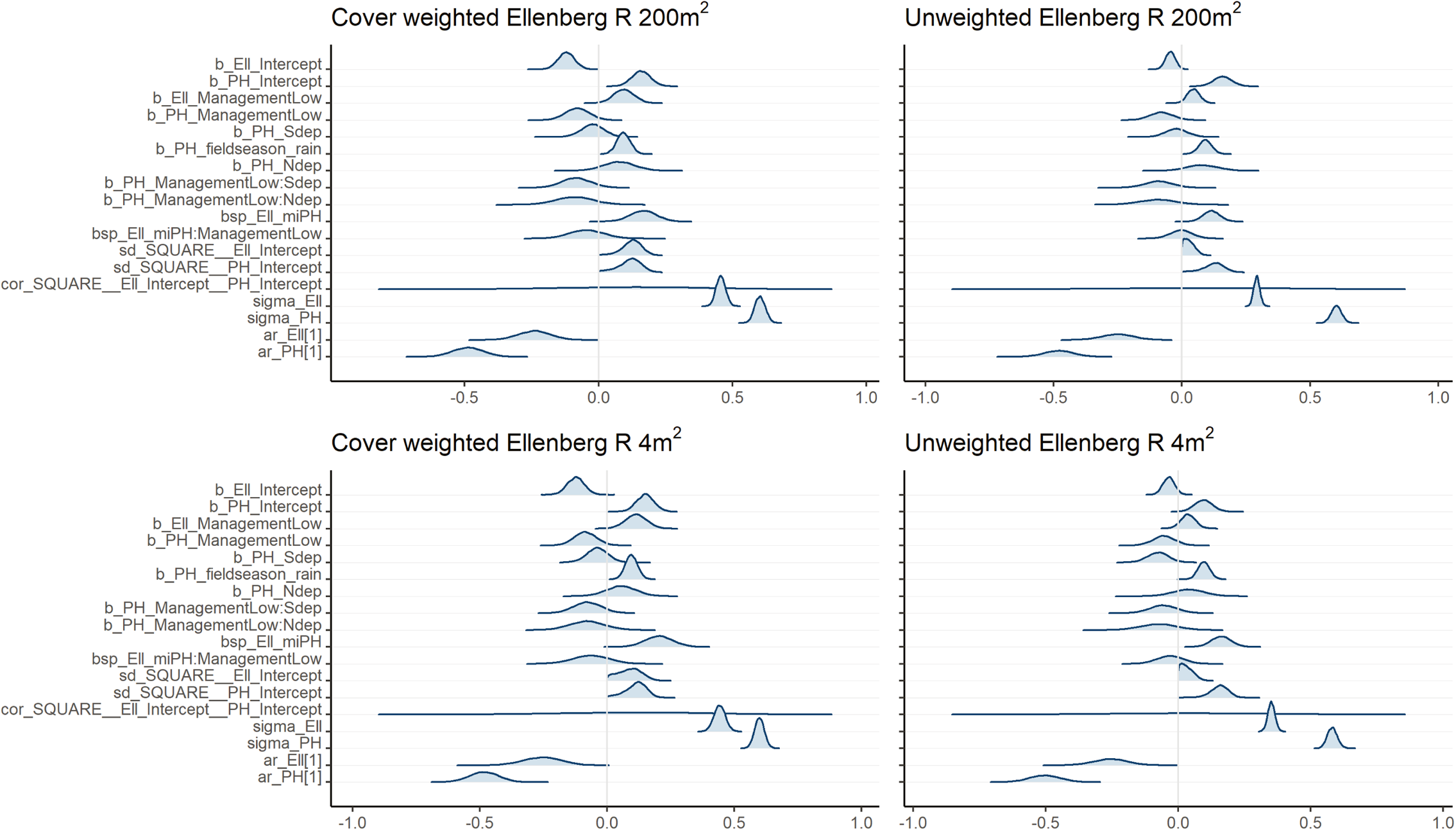
Measurement error multivariate models parameter estimates

**Figure 6:**
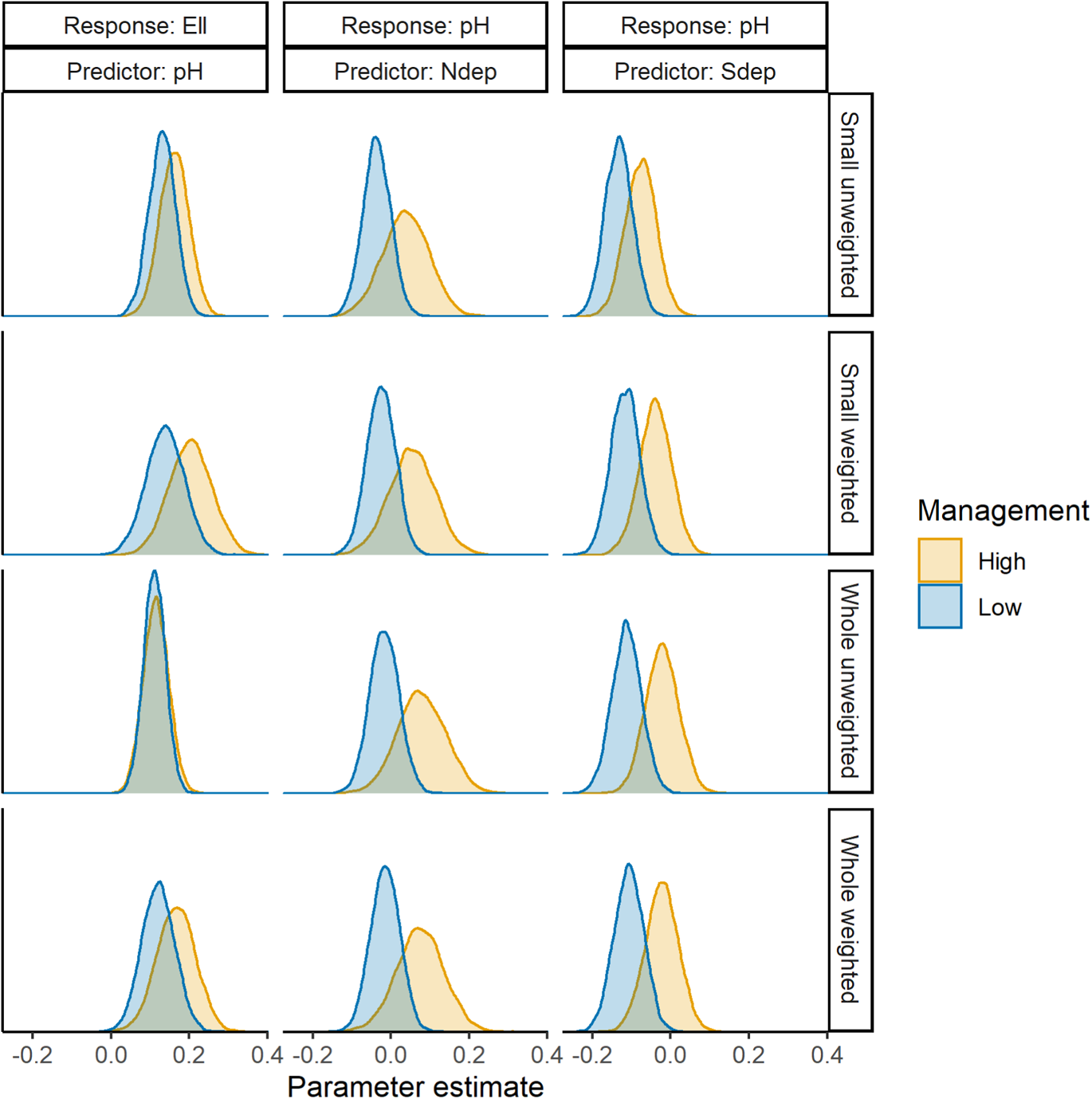
Parameter estimates for the management interaction parameters. Parameters are the estimated impact of one unit of change of soil pH, 72 units of change of N deposition and 4.5 units of change in S deposition due to the scaling by mean and standard deviation of N and S deposition.

**Figure 7:**
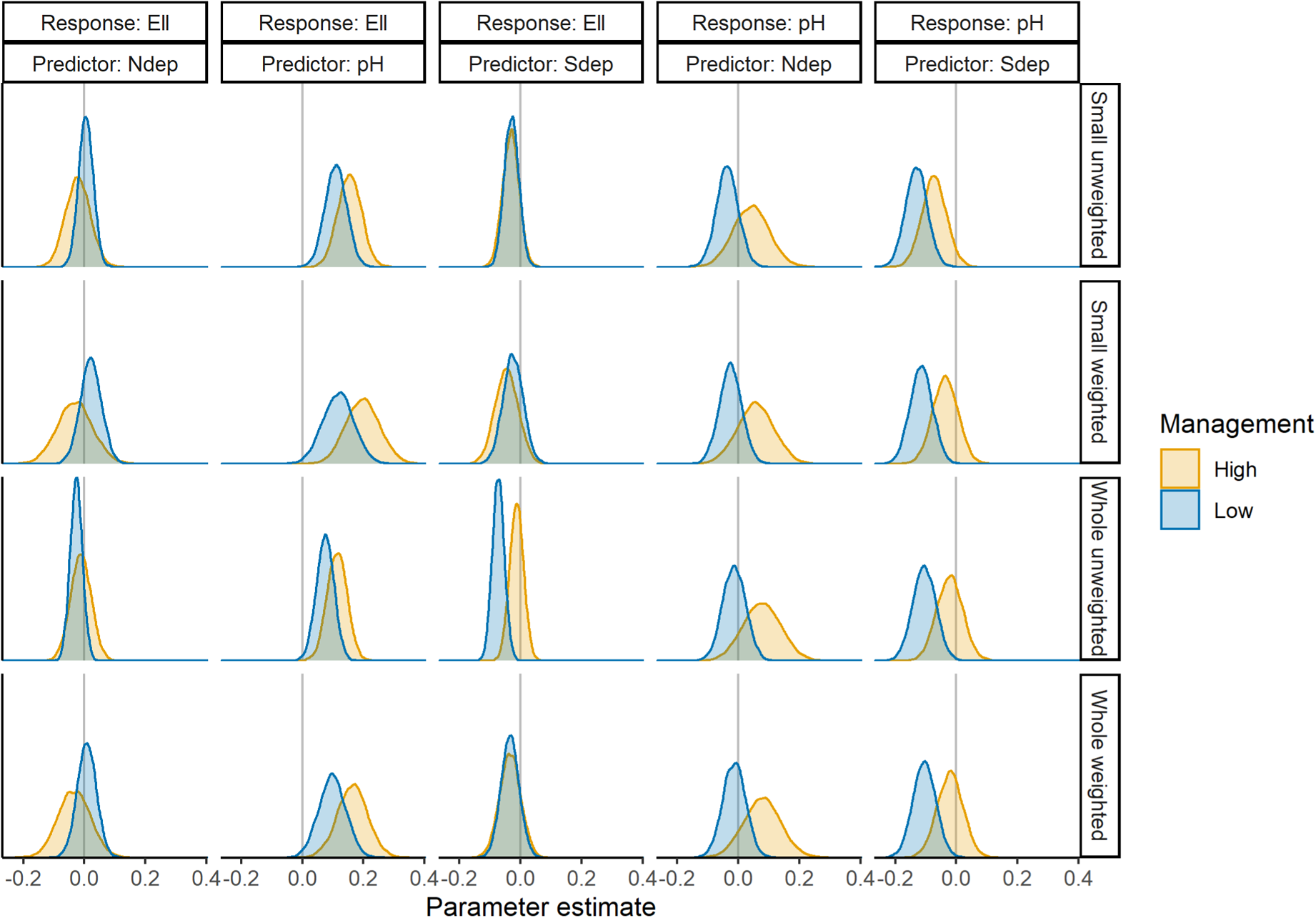
Parameter estimates of N deposition, S deposition and pH change effects upon Ellenberg R and soil pH change for the two management intensities.

## Supplementary Tables

**Table 1:**
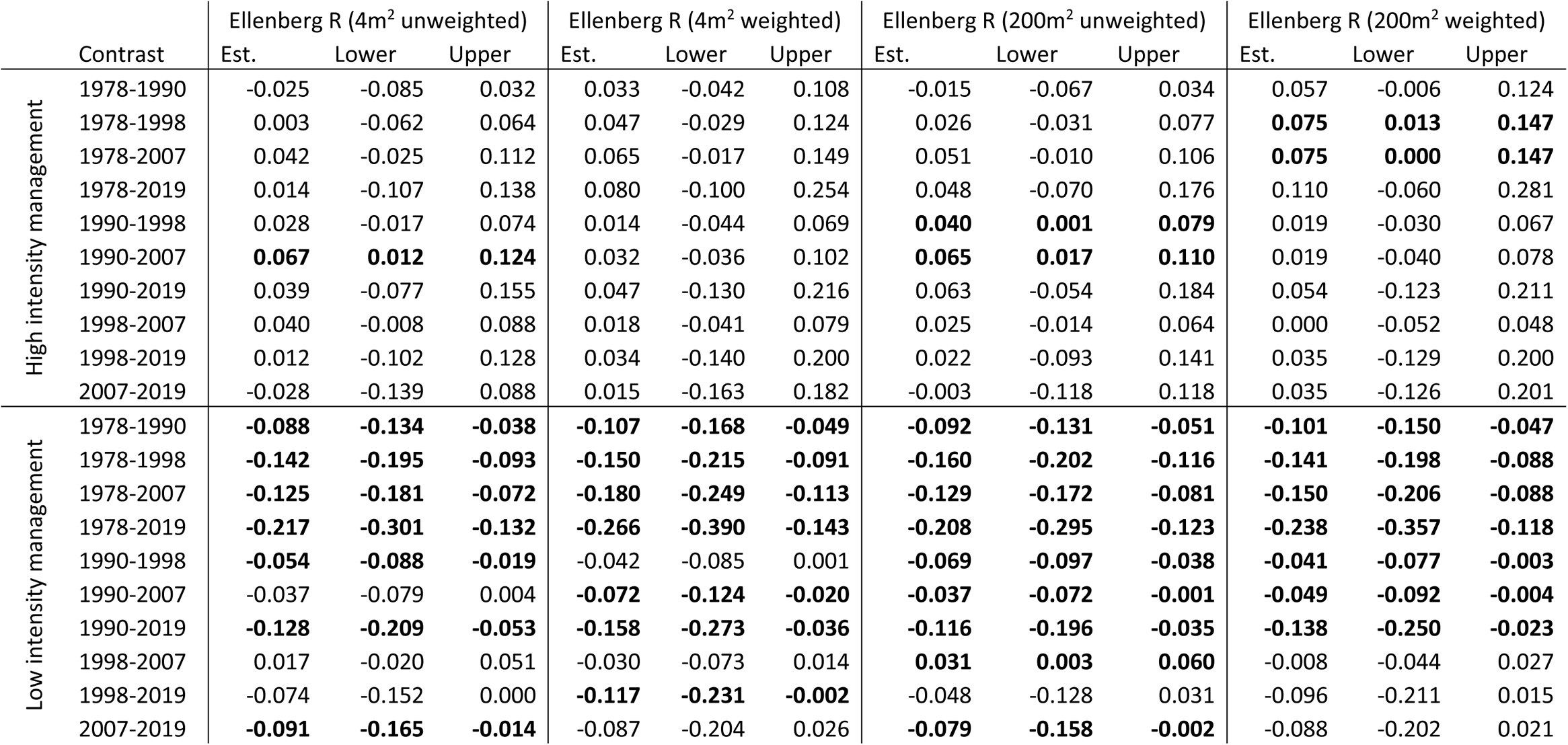
The estimated difference between years for mean Ellenberg for all measurements of Ellenberg R. The estimated difference and the lower and upper highest posterior density (95%) are given for each year contrast. Differences where the 95% HPD do not cross zero are in bold.

|  | Contrast | Ellenberg R (4m<sup>2</sup> unweighted) |  |  | Ellenberg R (4m<sup>2</sup> weighted) |  |  | Ellenberg R (200m<sup>2</sup> unweighted) |  |  | Ellenberg R (200m<sup>2</sup> weighted) |  |  |
| --- | --- | --- | --- | --- | --- | --- | --- | --- | --- | --- | --- | --- | --- |
|  |  | Est. | Lower | Upper | Est. | Lower | Upper | Est. | Lower | Upper | Est. | Lower | Upper |
| High intensity management | 1978-1990 | -0.025 | -0.085 | 0.032 | 0.033 | -0.042 | 0.108 | -0.015 | -0.067 | 0.034 | 0.057 | -0.006 | 0.124 |
|  | 1978-1998 | 0.003 | -0.062 | 0.064 | 0.047 | -0.029 | 0.124 | 0.026 | -0.031 | 0.077 | **0.075** | **0.013** | **0.147** |
|  | 1978-2007 | 0.042 | -0.025 | 0.112 | 0.065 | -0.017 | 0.149 | 0.051 | -0.010 | 0.106 | **0.075** | **0.000** | **0.147** |
|  | 1978-2019 | 0.014 | -0.107 | 0.138 | 0.080 | -0.100 | 0.254 | 0.048 | -0.070 | 0.176 | 0.110 | -0.060 | 0.281 |
|  | 1990-1998 | 0.028 | -0.017 | 0.074 | 0.014 | -0.044 | 0.069 | **0.040** | **0.001** | **0.079** | 0.019 | -0.030 | 0.067 |
|  | 1990-2007 | **0.067** | **0.012** | **0.124** | 0.032 | -0.036 | 0.102 | **0.065** | **0.017** | **0.110** | 0.019 | -0.040 | 0.078 |
|  | 1990-2019 | 0.039 | -0.077 | 0.155 | 0.047 | -0.130 | 0.216 | 0.063 | -0.054 | 0.184 | 0.054 | -0.123 | 0.211 |
|  | 1998-2007 | 0.040 | -0.008 | 0.088 | 0.018 | -0.041 | 0.079 | 0.025 | -0.014 | 0.064 | 0.000 | -0.052 | 0.048 |
|  | 1998-2019 | 0.012 | -0.102 | 0.128 | 0.034 | -0.140 | 0.200 | 0.022 | -0.093 | 0.141 | 0.035 | -0.129 | 0.200 |
|  | 2007-2019 | -0.028 | -0.139 | 0.088 | 0.015 | -0.163 | 0.182 | -0.003 | -0.118 | 0.118 | 0.035 | -0.126 | 0.201 |
| Low intensity management | 1978-1990 | **-0.088** | **-0.134** | **-0.038** | **-0.107** | **-0.168** | **-0.049** | **-0.092** | **-0.131** | **-0.051** | **-0.101** | **-0.150** | **-0.047** |
|  | 1978-1998 | **-0.142** | **-0.195** | **-0.093** | **-0.150** | **-0.215** | **-0.091** | **-0.160** | **-0.202** | **-0.116** | **-0.141** | **-0.198** | **-0.088** |
|  | 1978-2007 | **-0.125** | **-0.181** | **-0.072** | **-0.180** | **-0.249** | **-0.113** | **-0.129** | **-0.172** | **-0.081** | **-0.150** | **-0.206** | **-0.088** |
|  | 1978-2019 | **-0.217** | **-0.301** | **-0.132** | **-0.266** | **-0.390** | **-0.143** | **-0.208** | **-0.295** | **-0.123** | **-0.238** | **-0.357** | **-0.118** |
|  | 1990-1998 | **-0.054** | **-0.088** | **-0.019** | -0.042 | -0.085 | 0.001 | **-0.069** | **-0.097** | **-0.038** | **-0.041** | **-0.077** | **-0.003** |
|  | 1990-2007 | -0.037 | -0.079 | 0.004 | **-0.072** | **-0.124** | **-0.020** | **-0.037** | **-0.072** | **-0.001** | **-0.049** | **-0.092** | **-0.004** |
|  | 1990-2019 | **-0.128** | **-0.209** | **-0.053** | **-0.158** | **-0.273** | **-0.036** | **-0.116** | **-0.196** | **-0.035** | **-0.138** | **-0.250** | **-0.023** |
|  | 1998-2007 | 0.017 | -0.020 | 0.051 | -0.030 | -0.073 | 0.014 | **0.031** | **0.003** | **0.060** | -0.008 | -0.044 | 0.027 |
|  | 1998-2019 | -0.074 | -0.152 | 0.000 | **-0.117** | **-0.231** | **-0.002** | -0.048 | -0.128 | 0.031 | -0.096 | -0.211 | 0.015 |
|  | 2007-2019 | **-0.091** | **-0.165** | **-0.014** | -0.087 | -0.204 | 0.026 | **-0.079** | **-0.158** | **-0.002** | -0.088 | -0.202 | 0.021 |

**Table 2:** Parameter estimates for the four different measurement error models.

| Response | Parameter | Cover weighted 200m<sup>2</sup> |  | Unweighted 200m<sup>2</sup> |  | Cover weighted 4m<sup>2</sup> |  | Unweighted 4m<sup>2</sup> |  |
| --- | --- | --- | --- | --- | --- | --- | --- | --- | --- |
|  |  | Estimate | 95% quantile | Estimate | 95% quantile | Estimate | 95% quantile | Estimate | 95% quantile |
| EII | Intercept | -0.121 | -0.182 - -0.059 | -0.045 | -0.084 - -0.007 | -0.124 | -0.19 - -0.057 | -0.036 | -0.08 - 0.008 |
| EII | ManagementLow | 0.094 | 0.017 - 0.169 | 0.044 | -0.004 - 0.093 | 0.113 | 0.03 - 0.195 | 0.037 | -0.019 - 0.093 |
| EII | pH | 0.166 | 0.069 - 0.262 | 0.115 | 0.052 - 0.179 | 0.203 | 0.096 - 0.312 | 0.162 | 0.088 - 0.235 |
| EII | PH:ManagementLow | -0.046 | -0.169 - 0.079 | -0.004 | -0.087 - 0.079 | -0.065 | -0.21 - 0.073 | -0.031 | -0.129 - 0.066 |
| pH | Intercept | 0.158 | 0.089 - 0.228 | 0.158 | 0.087 - 0.228 | 0.150 | 0.082 - 0.218 | 0.095 | 0.029 - 0.162 |
| pH | ManagementLow | -0.080 | -0.167 - 0.009 | -0.081 | -0.168 - 0.010 | -0.086 | -0.172 - 0.001 | -0.055 | -0.139 - 0.028 |
| pH | ManagementLow:Sdep | -0.084 | -0.190 - 0.024 | -0.091 | -0.200 - 0.016 | -0.077 | -0.182 - 0.029 | -0.057 | -0.156 - 0.042 |
| pH | ManagementLow:Ndep | -0.090 | -0.227 - 0.051 | -0.095 | -0.234 - 0.044 | -0.079 | -0.213 - 0.056 | -0.075 | -0.207 - 0.056 |
| pH | Sdep | -0.023 | -0.107 - 0.059 | -0.022 | -0.104 - 0.062 | -0.039 | -0.122 - 0.043 | -0.077 | -0.156 - 0.003 |
| pH | fieldseason_rain | 0.093 | 0.044 - 0.141 | 0.091 | 0.042 - 0.139 | 0.095 | 0.048 - 0.143 | 0.097 | 0.053 - 0.142 |
| pH | Ndep | 0.075 | -0.047 - 0.193 | 0.078 | -0.042 - 0.199 | 0.054 | -0.065 - 0.172 | 0.039 | -0.080 - 0.156 |
| EII | SQUARE s.d. | 0.132 | 0.057 - 0.195 | 0.030 | 0.001 - 0.074 | 0.111 | 0.020 - 0.185 | 0.042 | 0.002 - 0.094 |
| pH | SQUARE s.d. | 0.364 | 0.291 - 0.441 | 0.364 | 0.291 - 0.444 | 0.342 | 0.271 - 0.418 | 0.352 | 0.288 - 0.420 |
|  | Cor SQUARE EII pH | 0.126 | -0.374 - 0.599 | 0.086 | -0.501 - 0.629 | 0.099 | -0.464 - 0.614 | 0.040 | -0.529 - 0.576 |
| EII | Sigma | 0.456 | 0.422 - 0.491 | 0.292 | 0.269 - 0.315 | 0.442 | 0.398 - 0.487 | 0.352 | 0.325 - 0.379 |
| pH | Sigma | 0.603 | 0.564 - 0.642 | 0.602 | 0.562 - 0.642 | 0.599 | 0.561 - 0.637 | 0.582 | 0.545 - 0.62 |
| pH | Temporal autocorrelation | -0.488 | -0.602 - -0.375 | -0.48 | -0.595 - -0.363 | -0.485 | -0.601 - -0.374 | -0.504 | -0.62 - -0.39 |
| EII | Temporal autocorrelation | -0.241 | -0.358 - -0.126 | -0.252 | -0.379 - -0.13 | -0.256 | -0.403 - -0.115 | -0.256 | -0.386 - -0.127 |

**Table 3:** Parameter estimates for the four different models without measurement error.

| Response | Parameter | Cover weighted 200m<sup>2</sup> |  | Unweighted 200m<sup>2</sup> |  | Cover weighted 4m<sup>2</sup> |  | Unweighted 4m<sup>2</sup> |  |
| --- | --- | --- | --- | --- | --- | --- | --- | --- | --- |
|  |  | Estimate | 95% quantile | Estimate | 95% quantile | Estimate | 95% quantile | Estimate | 95% quantile |
| EII | Intercept | -0.111 | -0.169 - -0.052 | -0.041 | -0.077 - -0.005 | -0.113 | -0.175 - -0.050 | -0.034 | -0.074 - 0.007 |
| EII | ManagementLow | 0.088 | 0.014 - 0.161 | 0.040 | -0.005 - 0.086 | 0.107 | 0.030 - 0.184 | 0.038 | -0.012 - 0.090 |
| EII | pH | 0.116 | 0.049 - 0.183 | 0.085 | 0.040 - 0.129 | 0.138 | 0.065 - 0.211 | 0.116 | 0.064 - 0.167 |
| EII | PH:ManagementLow | -0.026 | -0.113 - 0.061 | -0.005 | -0.062 - 0.054 | -0.033 | -0.128 - 0.064 | -0.026 | -0.094 - 0.041 |
| pH | Intercept | 0.17 | 0.101 - 0.239 | 0.170 | 0.100 - 0.239 | 0.164 | 0.096 - 0.232 | 0.116 | 0.051 - 0.181 |
| pH | ManagementLow | -0.074 | -0.163 - 0.010 | -0.073 | -0.160 - 0.014 | -0.081 | -0.165 - 0.005 | -0.056 | -0.138 - 0.024 |
| pH | ManagementLow:Sdep | -0.074 | -0.180 - 0.030 | -0.074 | -0.180 - 0.031 | -0.071 | -0.174 - 0.030 | -0.054 | -0.154 - 0.047 |
| pH | ManagementLow:Ndep | -0.075 | -0.208 - 0.058 | -0.075 | -0.209 - 0.059 | -0.068 | -0.200 - 0.062 | -0.065 | -0.194 - 0.063 |
| pH | Sdep | -0.017 | -0.099 - 0.065 | -0.017 | -0.101 - 0.064 | -0.029 | -0.108 - 0.051 | -0.062 | -0.139 - 0.013 |
| pH | fieldseason_rain | 0.091 | 0.042 - 0.141 | 0.091 | 0.042 - 0.140 | 0.092 | 0.044 - 0.141 | 0.097 | 0.053 - 0.143 |
| pH | Ndep | 0.070 | -0.047 - 0.184 | 0.070 | -0.047 - 0.188 | 0.052 | -0.061 - 0.165 | 0.038 | -0.076 - 0.149 |
| EII | SQUARE s.d. | 0.134 | 0.059 - 0.193 | 0.033 | 0.002 - 0.079 | 0.106 | 0.011 - 0.184 | 0.037 | 0.002 - 0.090 |
| pH | SQUARE s.d. | 0.128 | 0.034 - 0.196 | 0.132 | 0.043 - 0.197 | 0.126 | 0.032 - 0.192 | 0.157 | 0.089 - 0.217 |
|  | Cor SQUARE EII pH | 0.112 | -0.363 - 0.564 | 0.121 | -0.460 - 0.639 | 0.114 | -0.414 - 0.597 | 0.058 | -0.507 - 0.584 |
| EII | Sigma | 0.595 | 0.569 - 0.621 | 0.409 | 0.393 - 0.426 | 0.673 | 0.645 - 0.703 | 0.491 | 0.473 - 0.510 |
| pH | Sigma | 0.743 | 0.712 - 0.775 | 0.742 | 0.711 - 0.774 | 0.739 | 0.709 - 0.770 | 0.724 | 0.695 - 0.753 |
| pH | Temporal autocorrelation | -0.372 | -0.454 - -0.291 | -0.374 | -0.455 - -0.295 | -0.369 | -0.449 - -0.288 | -0.381 | -0.461 - -0.301 |
| EII | Temporal autocorrelation | -0.181 | -0.258 - -0.103 | -0.172 | -0.243 - -0.102 | -0.180 | -0.259 - -0.102 | -0.184 | -0.258 - -0.112 |

**Table 4:** Measurement error estimates for the different parameters

| Time period | Ellenberg R<br>(weighted 200m<sup>2</sup>) | Ellenberg R<br>(unweighted 200m<sup>2</sup>) | Ellenberg R<br>(weighted 4m<sup>2</sup>) | Ellenberg R<br>(unweighted 4m<sup>2</sup>) | pH (DIW) | pH (CaCl<sub>2</sub>) |
| --- | --- | --- | --- | --- | --- | --- |
| 1978-1998 | 0.382 | 0.284 | 0.503 | 0.333 | 0.461 | - |
| 1998-2007 | 0.397 | 0.308 | 0.562 | 0.385 | 0.446 | - |
| 2007-2019 | 0.526 | 0.297 | 0.672 | 0.443 | 0.303 | 0.257 |

